# Gap junctions in Turing-type periodic feather pattern formation

**DOI:** 10.1101/2023.04.15.537019

**Authors:** Chun-Chih Tseng, Thomas E. Woolley, Ting-Xin Jiang, Ping Wu, Philip K. Maini, Randall B. Widelitz, Cheng-Ming Chuong

## Abstract

Periodic patterning requires coordinated cell-cell interactions at the tissue level. Turing showed, using mathematical modeling, how spatial patterns could arise from the reactions of a diffusive activator-inhibitor pair in an initially homogenous two-dimensional field. Most activators and inhibitors studied in biological systems are proteins, and the roles of cell-cell interaction, ions, bioelectricity, etc. are only now being identified. Gap junctions (GJs) mediate direct exchanges of ions or small molecules between cells, enabling rapid long-distance communications in a cell collective. They are therefore good candidates for propagating non-protein-based patterning signals that may act according to the Turing principles. Here, we explore the possible roles of GJs in Turing-type patterning using feather pattern formation as a model. We found seven of the twelve investigated GJ isoforms are highly dynamically expressed in the developing chicken skin. *In ovo* functional perturbations of the GJ isoform, connexin 30, by siRNA and the dominant-negative mutant applied before placode development led to disrupted primary feather bud formation, including patches of smooth skin and buds of irregular sizes. Later, after the primary feather arrays were laid out, inhibition of gap junctional intercellular communication in the *ex vivo* skin explant culture allowed the emergence of new feather buds in temporal waves at specific spatial locations relative to the existing primary buds. The results suggest that gap junctional communication may facilitate the propagation of long-distance inhibitory signals. Thus, the removal of GJ activity would enable the emergence of new feather buds if the local environment is competent and the threshold to form buds is reached. We propose Turing-based computational simulations that can predict the appearance of these ectopic bud waves. Our models demonstrate how a Turing activator-inhibitor system can continue to generate patterns in the competent morphogenetic field when the level of intercellular communication at the tissue scale is modulated.

## Introduction

In 1952, Alan Turing published his seminal paper describing how spatial patterns can arise from two reactive and diffusive factors [1], termed morphogens, in a system which exhibits spatially uniform steady states that are stable in the absence of diffusion. However, 70 years later, we still have a limited understating of the identity of the morphogens and how these morphogens can propagate through complex biological systems [2, 3]. Here we use the developing chicken skin to further our understanding. During embryonic development, feathers form in discrete regions over the body surface, called tracts or pterylae, that are distributed across the avian skin (Figure 1A) [4]. Featherless regions lying between tracts are referred to as apteric regions. Chicken skin exhibits characteristic hexagonal feather arrays, which are best demonstrated in the spinal tract. In the spinal tract, the skin forms a competent feather field and then the first row of feather primordia appears along the midline. Subsequent rows of feathers spread bilaterally, adding more rows of younger feather primordia (Figure 1B) [5, 6]. These have been documented to form due to chemical reactions and mechanical forces that trigger the self-organization of mesodermal cells. The resultant mesodermal cell aggregates then promote the epidermal expression and nucleation of a key regulator of feather morphogenesis, β–catenin [7, 8]. The periodicity of the feather primordia is at least in part ensured by a global ectodysplasin A (EDA) patterning wave expanding from the midline of the spinal tract (Figure 1B), which lowers the mesenchymal cell density threshold needed to form feather primordia [9, 10]. Locally, Turing’s theory of diffusion-driven instability describes how a pair of reaction-diffusion (RD) equations with activator-inhibitor kinetics and randomly perturbed initial conditions in a homogeneous field can lead to periodic pattern formation [1]. Based on this model, a self-amplifying short-range activator can stimulate the production of a long-range inhibitor that, in turn, negatively regulates the activator. Previous studies showed that a variety of spatial patterns can be generated in simulated mathematical RD model systems [3, 11, 12]. In addition, a global patterning wave involving mechanochemical reactions allows the dermal cells to aggregate [9, 13, 14] and together they produce a highly ordered hexagonal patterned array of feather primordia in the chicken. On the other hand, without the global wave, local Turing patterning can produce periodic buds simultaneously in the tract [9, 14], similar to the patterning process in reconstituted feather explants [7].

**Figure 1.**
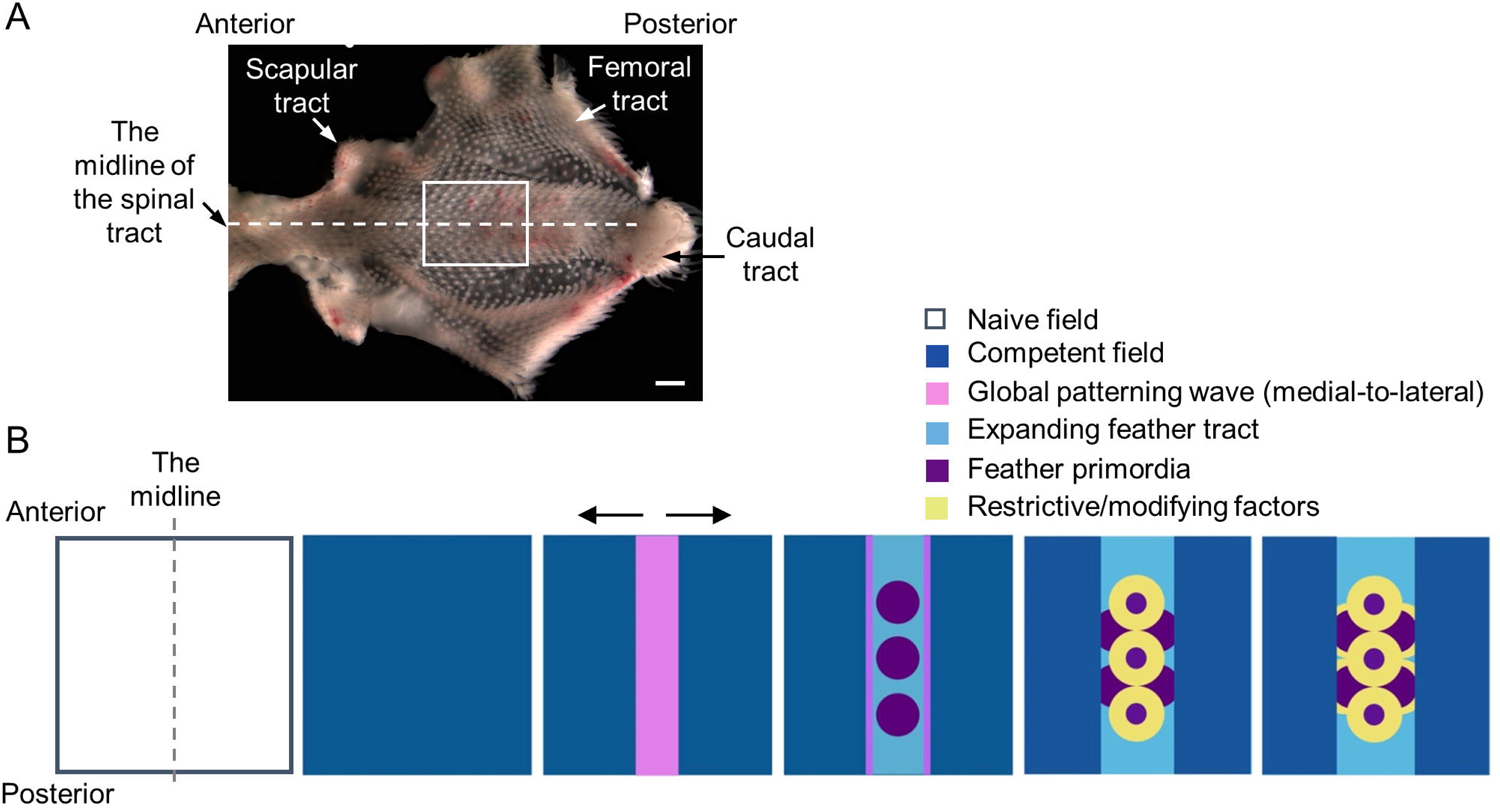
Schematic illustrating the emergence of chicken feather primordia. (A) The chick skin contains discrete feathered areas, termed tracts. In E14.5 chick dorsal skin we highlight four of the tracts: the spinal tract, the scapular tract, the femoral tract, and the caudal tract. Anterior of the embryo is toward the left. Scale bar, 1000 µm. Developmental changes that occur within the boxed region of the thoracic-lumbar region are schematized below. (B) Feather formation starts from a field lacking any evident patterning cues, followed by the emergence of competent patterning fields. In the spinal tract, the highly ordered feather array is formed along the midline and then progressively and bi-laterally expanded across the skin. While the patterning wave(s) travel through the skin, the spatiotemporal pattern and the size of feather primordia are sequentially established by promoting and restricting/modifying factors. The triangle-headed arrows show the directions of the patterning waves. Anterior is to the upper side.

Importantly, Turing’s RD theory can be applied to a diverse range of integuments, such as the FGF/SHH-BMP pair in periodic feather patterning, shark skin denticle patterning [9, 15–17] and the WNT-DKK pair in hair follicle spacing [18]. Most of the morphogens discovered in animals are secreted proteins which diffuse. For example, *Pax6* establishes a *Tgfb2* and *Fst* Turing signaling network that has been proposed to pattern chick eye cup development [19]. The role of FGF, hedgehog, Wnt and BMP signaling in establishing the murine periodic striped patterning of rugae ridges on the oral palate was identified using inhibitors [20]. The roles of ions, bioelectricity, small molecules, and cell-cell contacts in periodic patterning are just emerging. Notably, cell-cell interactions between pigment cells are required for zebrafish stripe and spot patterning [21]. Although the underlying molecular mechanism of zebrafish pigment pattern formation is still elusive, it is, at least in part, regulated by spermine, a small poly-cation, and its regulation of the flow of plasma membrane channels, including the potassium channel, Kir7.1, and gap junctions, Cx41.8 and Cx39.4 [21–26].

Gap junctions (GJs) are formed by direct docking of connexon or pannexon hemichannels consisting of 6 subunits of connexin (Cx) or pannexin (Panx) family proteins, respectively, between adjacent cells. GJs allow the direct exchange of cellular content, including ions, small metabolites and second messengers, between neighboring cells [27–29]. Connexins and pannexins form distinct families of gap junction subunits. There are 21 connexins and 3 pannexins within the human genome and at least 12 connexins and 3 pannexins identified or predicted within the chick genome. The conserved domains, including N-terminus, extracellular loops and transmembrane domains in connexin families, have shown great sequence homology across vertebrate species [30]. Pannexins share some sequence homology with invertebrate gap junctions, innexins, but show no sequence homology with connexins [31]. GJs formed by Cx or Panx possess distinct electrophysiological and pharmacological properties. Panx GJs are less sensitive to the change in trans-junctional voltage compared to Cx GJs, and Panx GJs favor anionic substrates over cations [32]. Many of these GJ isoforms exhibit tissue specific and overlapping expression patterns. The “knock in” experiment, a genetic technique that replaces a gene with another one in the genomic locus, performed in mouse showed that individual connexins may play distinct and shared roles in specific cellular processes [33]. In agreement with this, studies have found that the heterotypic and heteromeric types of connexins are able to form functional channels. GJs formed by different types of Cxs exhibited a selective permeability to cytoplasmic molecules [34]. Importantly, many studies have demonstrated that GJs not only mediate the intercellular exchange of hydrophilic molecules but also act as a regulatable signaling scaffold that can be finely tuned by posttranslational modifications, such as phosphorylation and ubiquitination and form complexes with a variety of junctional and signaling molecules [34–36].

Connexins and pannexins are expressed in diverse patterns in skin and skin appendages [37–40]. Transcriptome profiling has revealed that multiple connexin and pannexin isoforms are expressed during embryonic feather morphogenesis (H&H stages 31 and 35). Among them, Cx43 is the most abundantly expressed isoform [38]. The same study examined the functions of Cx43 in greater detail and showed that Cx43 can facilitate the propagation of intercellular calcium signaling and regulate mesenchymal cell migration during elongation of feather buds [38]. In an earlier study, functional coupling of gap junctions between cells during early feather morphogenesis was demonstrated by the transfer of lucifer yellow (LY) injected into a single cell, where the gap junction intracellular communication (GJIC) exhibits asymmetric diffusion and differential compartmentalization between bud and interbud regions in the chick skin (H&H stage 30 to 35) [41].

Periodic feather pattern formation requires coordinated cell-cell interactions. The properties of GJs that allow them to facilitate the exchange of cellular information between adjacent cells and acts as a signaling platform at cell-cell junctions make the gap junctions a viable candidate to be involved in feather patterning. The previous studies mentioned above have shown that GJIC is a highly dynamic process and is associated with compartmentalization during feather morphogenesis. GJIC and the specific GJ isoform, Cx43, are important for the elongation of individual feather buds. However, these studies only addressed limited aspects of the functional importance of GJIC during feather morphogenesis, and the roles of GJ isoforms in periodic patterning are still largely unknown. In the present study, we focus on the connexin family and show that seven out of twelve tested connexins are expressed during early feather morphogenesis (H&H stage 28 to 35). Suppression of GJIC activities by small molecule gap junction channel blockers, 18 α-glycyrrhetinic acid (AGA) and its derivative, in *ex vivo* cultured chicken skin explants allows the emergence of ectopic feather buds through consecutive Turing instabilities or in specific locally competent tissues. AGA treatment activates the expression of *β-catenin* and *Shh*, early markers of feather morphogenesis, and stimulates cell proliferation in the ectopic buds. These results suggest that diffusing factors passing through gap junctions during normal feather patterning may serve as the long-range inhibitory signals described in Turing’s model.

Additionally, we found that Cx30 is the earliest expressed connexin isoform during feather patterning as demonstrated by whole-mount *in situ* hybridization. Functional perturbations of Cx30 by introducing siRNA and RCAS virus carrying a COOH-terminal truncated Cx30 mutant (a.a. 1-214) resulted in dysregulated development and/or patterns of feather primordia. It is worthwhile to mention that Clouston syndrome, a human disease resulting from Cx30 mutations and showing hair loss and skin abnormalities, exhibits similar feather loss phenotypes as those observed in the Cx30-perturbed experiments. Overall, this study uncovers unexpected roles of GJs in feather patterning and provides new insights into how long-range direct diffusion through GJs can be coupled with Turing’s mechanism and local tissue competence, referred to the ability of tissues to sense and respond to environmental stimuli, to modulate diverse patterning outcomes at different morphogenetic stages.

## Results

### Connexins exhibit dynamic expression patterns during early feather morphogenesis

We used the identified or predicted NCBI nucleotide coding sequence for 12 chicken connexin isoforms to examine their expression at early stages (H&H stage 28 to 35, corresponding to E6 to E9) of feather morphogenesis using whole-mount *in situ* hybridization (WM-ISH) and tissue sectioning of the stained embryos. To avoid cross reactivity between connexin isoforms, we designed primer sets for each connexin ensuring that WM-ISH probe sequences shared less than 55% similarity (S1 Table).

Connexin 30 (Cx30) exhibited highly dynamic expression patterns at the pre-placode stage (H&H stage 28, St28) to the short bud stage (H&H stage 34) but was not detected at the long bud stage (H&H stage 35) (Figure 2A). At H&H stage 28, it emerged in the epithelium of the spinal tract as a midline stripe and then gradually became restricted to the bud epithelium at H&H stage 31 (Figure 2A and B). During feather primordia stabilization and polarization, Cx30 expression in the bud became further restricted in size at the short bud stage anterior epithelium. Restricted Cx30 expression corresponds to the expanding inhibitory zones as marked by the dashed circles in Figure 2B. The emerging Cx30 transcripts also were coupled with bi-lateral global traveling wave expansion, highlighted by arrows pointing to the partial circle/triangular shaped *in situ* hybridization pattern (Figure 2B). The early Cx30 expression prompted us to compare its expression with β–catenin, the feather tract and early placode marker. We then compared it with Cx43, a transcriptional β–catenin target that was previously identified by bulk RNA-seq as the most abundantly expressed GJ during early feather patterning [38]. The transitional bud morphogenesis process can be best observed in the femoral tract. By independently staining contralateral halves of embryos, we can compare expression patterns of two different genes in the same embryo. As shown in Figure 2C, we observed the same number of rows of β–catenin and Cx30 expression, but β–catenin showed expression in one additional primordium in row 7 compared to Cx30 (arrows). On the other hand, Cx30 expression preceded Cx43 expression by one additional row. Thus, we can conclude that among these three genes, β–catenin is expressed earliest followed by Cx30 and then Cx43. Interestingly, it appears that Cx30 and Cx43 expression were mutually exclusive in the bud domain in more mature feather primordia. This can easily be seen in the fourth feather row of embryo 2, where Cx43 is expressed as a hollow circle while Cx30 is expressed as a central dot in feather primordia localized at comparable sites (Figure 2C). A similar hollow Cx43 pattern could also be observed in more mature feather primordia within the spinal tract (S1 Figure).

**Figure 2.**
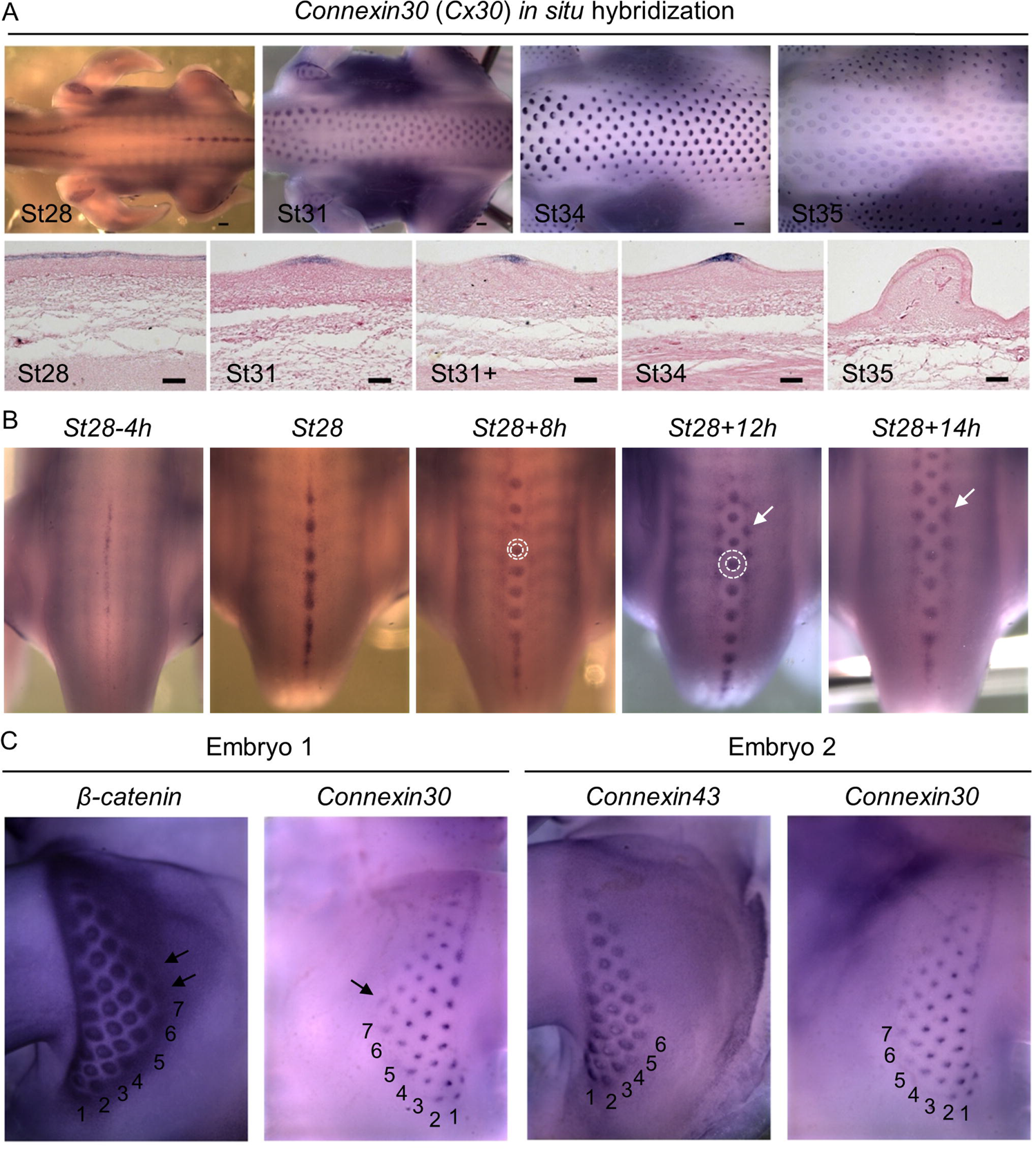
Dynamic expression of connexin 30 (Cx30) during early feather morphogenesis. (A) Cx30 expression in H&H stage (St) 28 to 35 chicken embryos. Upper panels: Cx30 RNA was visualized by whole-mount in situ hybridization (WM-ISH). Lower panels: After WM-ISH, the embryos were embedded in paraffin and then longitudinally sectioned along or near the midline of the spinal tract. Tissue sections were counter-stained with 10% eosin Y. Thickness of tissue sections: 14 µm. Anterior is to the left. Scale bars, 300 µm (whole embryo); 50 µm (tissue sections). (B) Cx30 RNA was visualized by WM-ISH in chicken embryos at or around H&H stage 28. Left to right: embryos were collected from earlier to later time points as indicated. Dashed circles highlight feather primordia and the adjacent regions. The triangle-headed arrows indicate emerging feather primordia. (h, hours). Anterior is toward the top. (C) Gene expression of β–catenin, Cx30, or Cx43 in the femoral tract was visualized by WM-ISH on contralateral sides of embryos at H&H stage 31. The numbers indicate feather primordium rows. The triangle-headed arrows, emerging feather primordia. Anterior of the embryo is toward the top.

Connexin 43 showed highly dynamic expression patterns during feather patterning at H&H stage 28 to 35 as demonstrated by WM-ISH and tissue sectioning of the stained embryos (Figure 3A). At the pre-placode stage (H&H stage 28), Cx43 is weakly expressed in both the mesoderm and the epithelium in the entire spinal tract with enhanced expression in the midline (solid arrow). Since the resolution of WM-ISH could not clearly demonstrate whether Cx43 expression is localized to the midline at St28 (Figures 3A and 3B, upper panels), we stained embryos with a chick Cx43 antibody (Santa Cruz Biotechnology, sc-9059) and showed that it was present in the entire spinal tract skin, exhibiting enhanced midline expression (Figure 3B, lower panels). Moreover, Cx43 protein was expressed not only at pre-placode stage (H&H stage28) but also at a much earlier stage (St28-24h). During placode formation (H&H stage 31), Cx43 expression was increased in the bud epithelium and undetectable in the mesoderm (Figure 3A and B). Interestingly, its expression swiftly became restricted to the interbud epithelium and mesoderm at H&H stage 34 when feather buds just started to show visible signs of asymmetric growth (Figure 3A). When the feather buds further elongate, Cx43 expression appears in the distal epithelium and mesoderm of the bud domain (Fig 3A, St34+) and extends to the proximal region of the bud mesoderm (St35). Cx43 expression is absent in the interbud at St34+ and St35.

**Figure 3.**
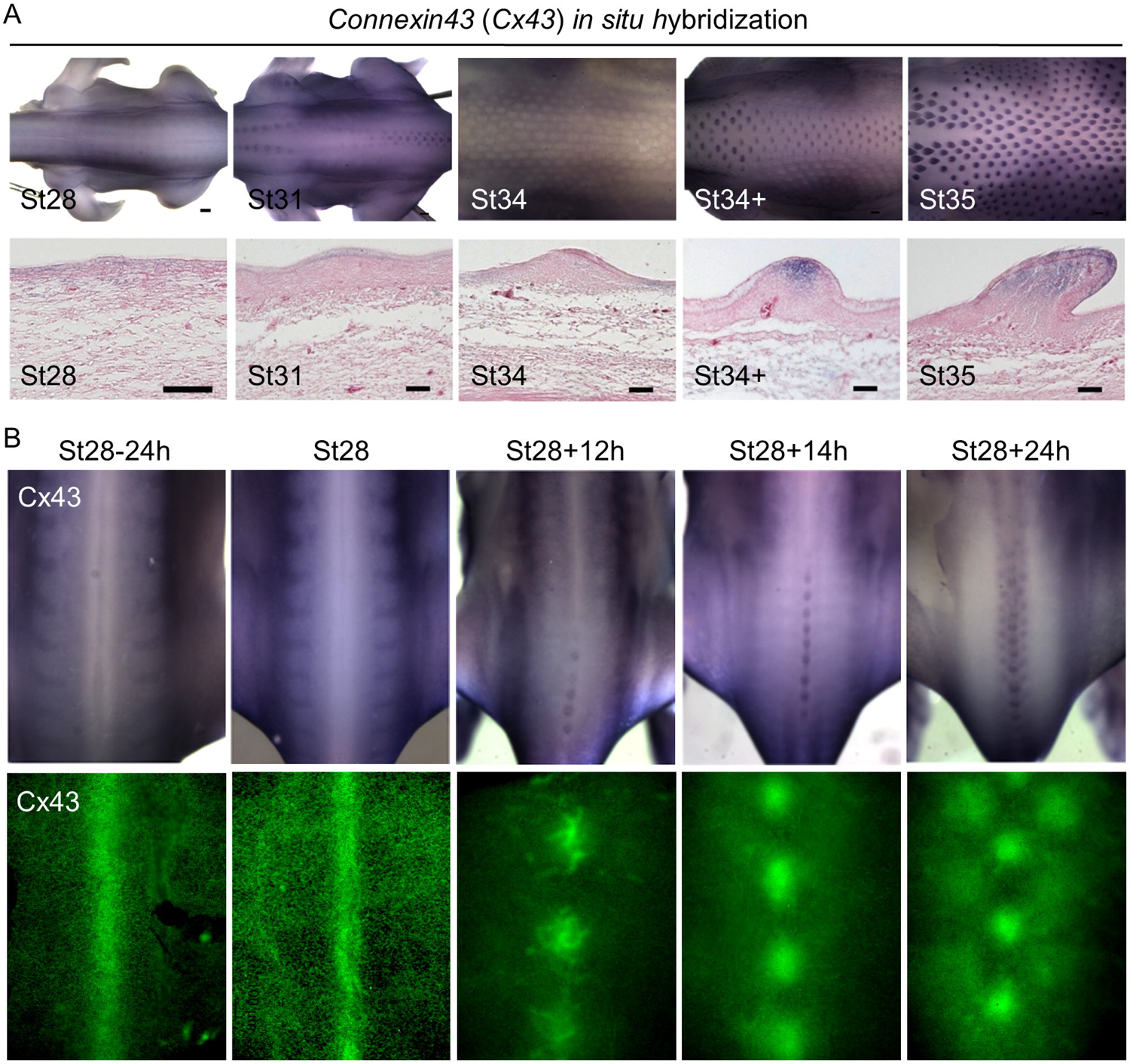
Expression Cx43 during early feather morphogenesis. (A) Gene expression of Cx43 in the chicken embryos between H&H stage 28 and 35. Upper panels: Cx43 RNA was visualized by WM-ISH. Lower panels: After WM-ISH, the embryos were embedded in paraffin and then longitudinally sectioned along or near the midline of the spinal tract. Tissue sections were counter-stained with 10% eosin Y. Thickness of tissue sections: 14 µm. Anterior is to the left. Scale bars, 300 µm (whole embryo); 50 µm (tissue sections). (B) Upper panels: Cx43 RNA was visualized by WM-ISH in chicken embryos at, or around, H&H stage 28. Left to right: embryos were collected from an earlier to later time points. (h, hours). Lower panels: protein expression of Cx43 in the dorsal skin of chicken embryos at the corresponding developmental stages as the upper panels. The embryos were stained with an antibody against Cx43 (Santa Cruz Biotechnology, sc-9059) by WM-immunostaining and then the dorsal skins were surgically removed for imaging. Anterior is toward the top.

Connexin 40 exhibited dotted patterns in the epithelium at H&H stage 31 to 35 and was undetected at earlier stages (Figure 4A). At H&H stage 31, Cx40 was evenly expressed and then disappeared in the bud domain. From H&H stage 34 to 35, Cx40 expression was enhanced in the proximal posterior end and gradually covered the entire feather bud epithelium (Figure 4A). This transitional expression pattern could be best observed in the femoral tract (Figure 4A, middle panel). The dotted patterns suggest that Cx40 was associated with specific cellular activities or cell types. Indeed, studies have shown that connexin 41.8, an ortholog of chicken Cx40 in the zebrafish, could regulate stripe/spot patterning [42, 43]. In Japanese quails, melanocyte Cx40 expression could modulate body pigment stripe formation [44]. These studies strongly suggest that Cx40 was expressed in chick embryo precursor pigment cells.

**Figure 4.**
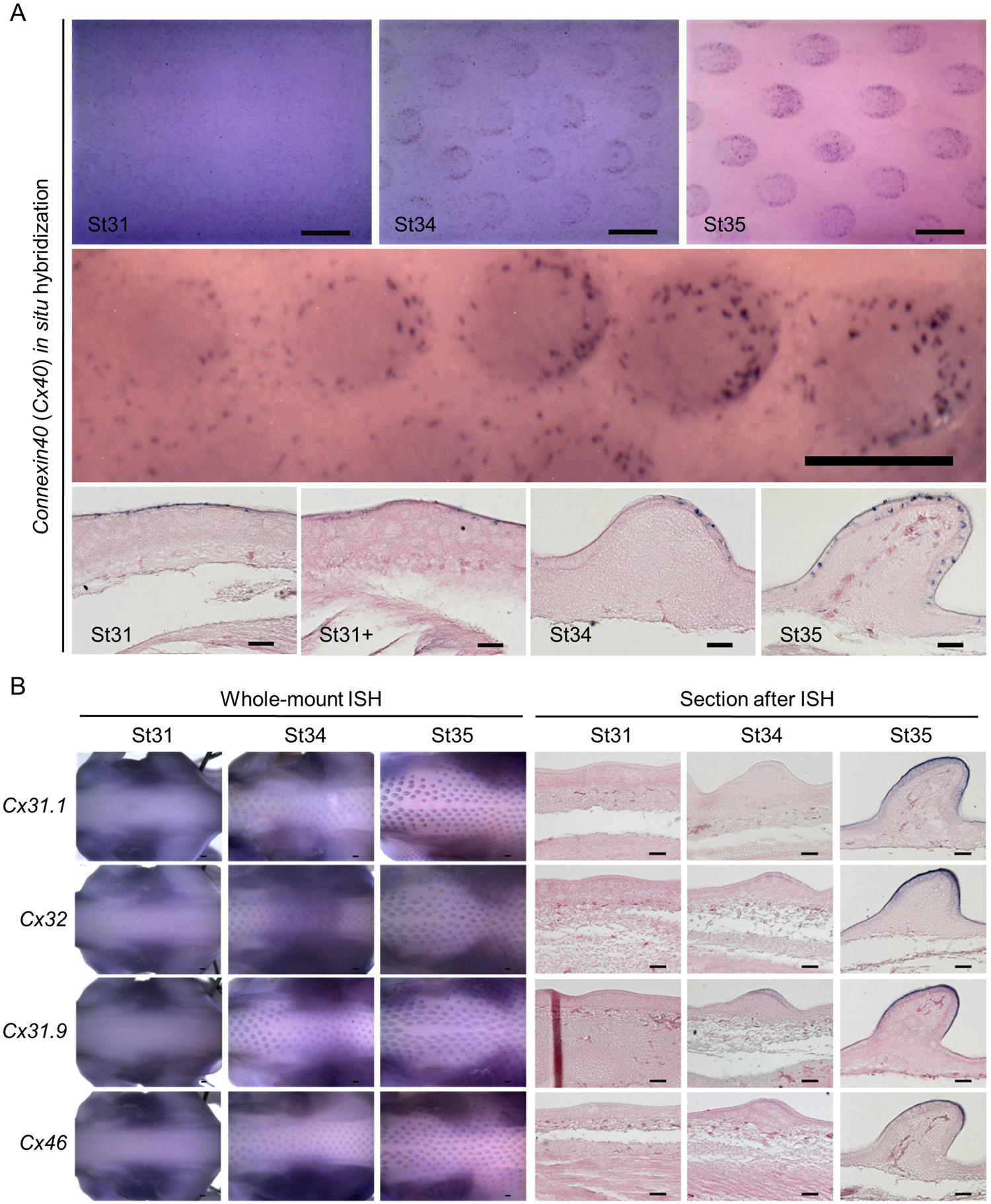
Expression of Cx40, Cx31.1, Cx32, Cx31.9, and Cx46 during early feather morphogenesis. (A) Gene expression of Cx40 in chicken embryos between H&H stage 31 and 35. Upper panels: dorsal views of embryonic thoracic-lumbar regions. Cx40 RNA was visualized by WM-ISH. The middle panel: Cx40 expression in the left embryonic femoral tract at H&H stage 35, which represents the development of feather buds in the midline of the spinal tract between H&H stage 31+ and 34. Lower panels: after WM-ISH, the embryos were embedded in paraffin and then longitudinally sectioned along or near the midline of the spinal tract. Tissue sections were counter-stained with 10% eosin Y. Thickness of tissue sections: 20 µm. Anterior is to the left. Scale bars, 300 µm (whole embryo); 50 µm (tissue sections). (B) Expression of the listed connexin isoforms shown by WM-ISH (left panels) and longitudinal sections along or near the midline of the chicken embryo spinal tract (right panels) at the stages between H&H stage 31 and 35. Anterior is to the left. Scale bars, 300 µm (whole embryo); 50 µm (tissue sections).

Other connexin isoforms, including Cx31.1, Cx32, Cx31.9 and Cx46, were detected by WM-ISH at later stages (Figure 4B). The expressions of Cx36, Cx37, Cx40.1, Cx50, and Cx52.6 were undetectable (data not shown). Both the Cx31.1 and Cx32 were weakly expressed, if expressed at all, at H&H stage 34 and strongly expressed in both bud and interbud epithelium at H&H stage 35. Cx31.9 expression could be identified at H&H stage 31 and afterwards in the outer bud epithelium. In the bud epithelium, Cx46 was weakly expressed at H&H stage 34 and more strongly expressed at H&H stage 35. These differential expression patterns in different bud and interbud epithelium layers suggest the roles that these connexins might play in epidermal differentiation and compartmental development. The schematic illustration in Figure 5 summarizes the expression patterns of all the connexins we studied, viewed from above the embryos (Figure 5A) or observed in tissue sections (Figure 5B). Collectively, these dynamic connexin expression patterns during feather morphogenesis suggest that they are active participants in early feather patterning. The distinct and overlapping patterns indicate connexins may have unique and shared functions during feather morphogenesis.

**Figure 5.**
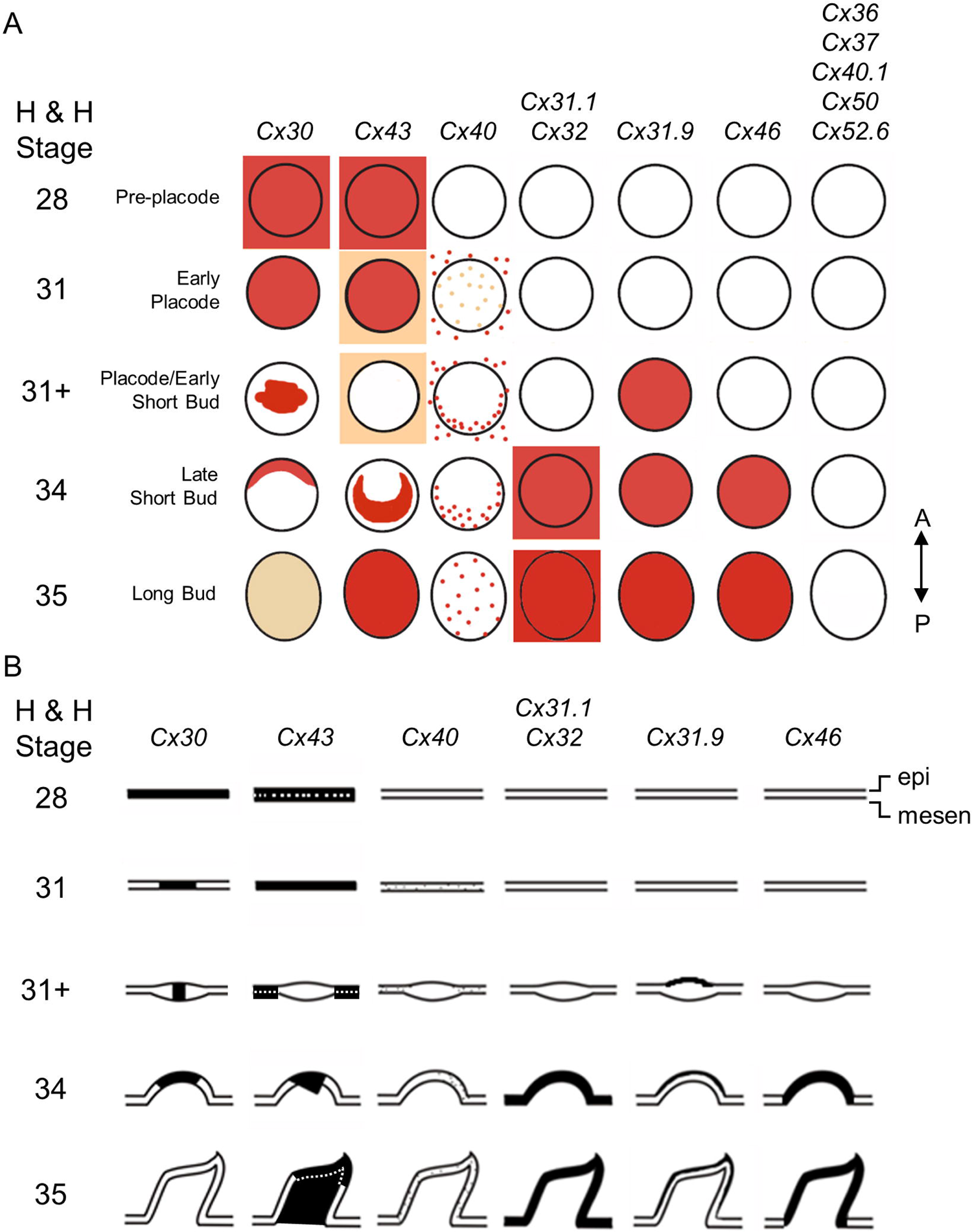
Schematic summary of connexin expression patterns during feather bud development. (A) Viewed from the top. Cx30 was expressed early in the emerging feather tract at H&H stage 28 and later (H&H stage 31) became restricted to the bud primordia. The expression gradually disappeared in the marginal regions of buds (H&H stage 31+) and eventually appeared in anterior bud regions (H&H stage 34). Expression was nearly undetectable at long bud stage (H&H stage 35). Cx43 was expressed in the feather-forming field (H&H stage 28) and gradually increased in the bud domain (H&H stage 31) but was completely gone at the early short bud stage (H&H stage 31+). At H&H stage 34, expression in the interbud domain disappeared while expression became intensified at more posterior side of the buds. Then (H&H stage 35) the signal occupied the entire bud domain. The remaining Cxs did not appear at initial stages (H&H stage 28). Among them, Cx40 shows a stippled pattern around the entire epithelium at St 31. This expression disappears transiently within the bud but gradually reappears in the posterior bud domain. At approximately the same time, the expression in interbuds was markedly lost. At H&H stage 35, Cx40 was expressed in entire bud domain with only sporadic expression detectable in the interbud. Cx31.1 and Cx32 were expressed with similar patterns in the bud and interbud domains starting from H&H stage 34 and beyond. Cx31.9 and Cx46 were expressed in bud domains starting from H&H stage 31+ and 34, respectively. Cx36, Cx37, Cx40.1, Cx50, and Cx52.6 were undetectable between H&H stage 28 and 35. A, anterior. P, posterior. (B) The schematic diagrams show the dynamic connexin expression in the sections after WM-ISH. Anterior is to the left. epi, epithelium. mesen, mesenchyme.

### Functional coupling of gap junctions during early feather patterning

Next, we investigated if gap junctions are functionally coupled between adjacent cells during early embryonic feather morphogenesis. Previously, our lab visualized GJIC in embryonic chicken skins by injecting lucifer yellow (LY) into a single cell and observed efficient LY dye transfer between cells in the H&H stage 30 to 32 embryonic placode epithelium [41]. In the same study, our lab also observed efficient LY dye coupling in the mesoderm but less so in the epithelium of short buds (equivalent to H&H stage 35 in this study). Dye transfer in the interbud epithelium at this stage is moderately efficient. Notably, LY dye transfer does not cross the bud epithelium and mesoderm boundary, nor the boundary of bud and interbud regions at the short bud stage. Here, we expanded this study and utilized previously described [45] scrape-loading of LY to visualize macro-scale GJIC in the spinal tract during early feather morphogenesis in the skin explants (Figure 6 and 7, H&H stage 28 to 35). We did not differentiate the localizations of LY dye transfer occurring in the epithelium or the mesoderm when we performed epifluorescence imaging due to the limitations of our imaging settings. We found that at pre-placode H&H stage 28, LY dye transfer (in green) efficiently crosses the dorsal skin (Figure 6A). Rhodamine dextran (Rho; 10 kDa, in red) is too big to pass through GJs and served as a control for the wound caused by the scrape-loading procedure. The intensities (grey values) of LY and Rho along the yellow lines (L0-L3) were measured with ImageJ software and used to create an X-Y scatter plot in Microsoft Excel. The yellow line L3 is a control for the background levels of the fluorescent intensities. The traveling distance (ΔD) indicates the distance the LY dye diffused away from the wound. Raw data of the measurement included in the dye coupling studies are shown in S1 Data. The results showed that LY dye transfer is faster at the anterior side (ante) than the posterior side (post) of the wound along the anterior-posterior (A-P) axis, suggesting that there is a preferred directionality toward the anterior side for LY dye transfer on the dorsal skin of chick H&H stage 28 embryos. The mechanisms underlying this phenomenon and how this may affect the macro-scale skin patterning need further research. We also observed a slightly faster LY dye transfer along the midline of the spinal tract (L0) compared to the adjacent region (L1).

**Figure 6.**
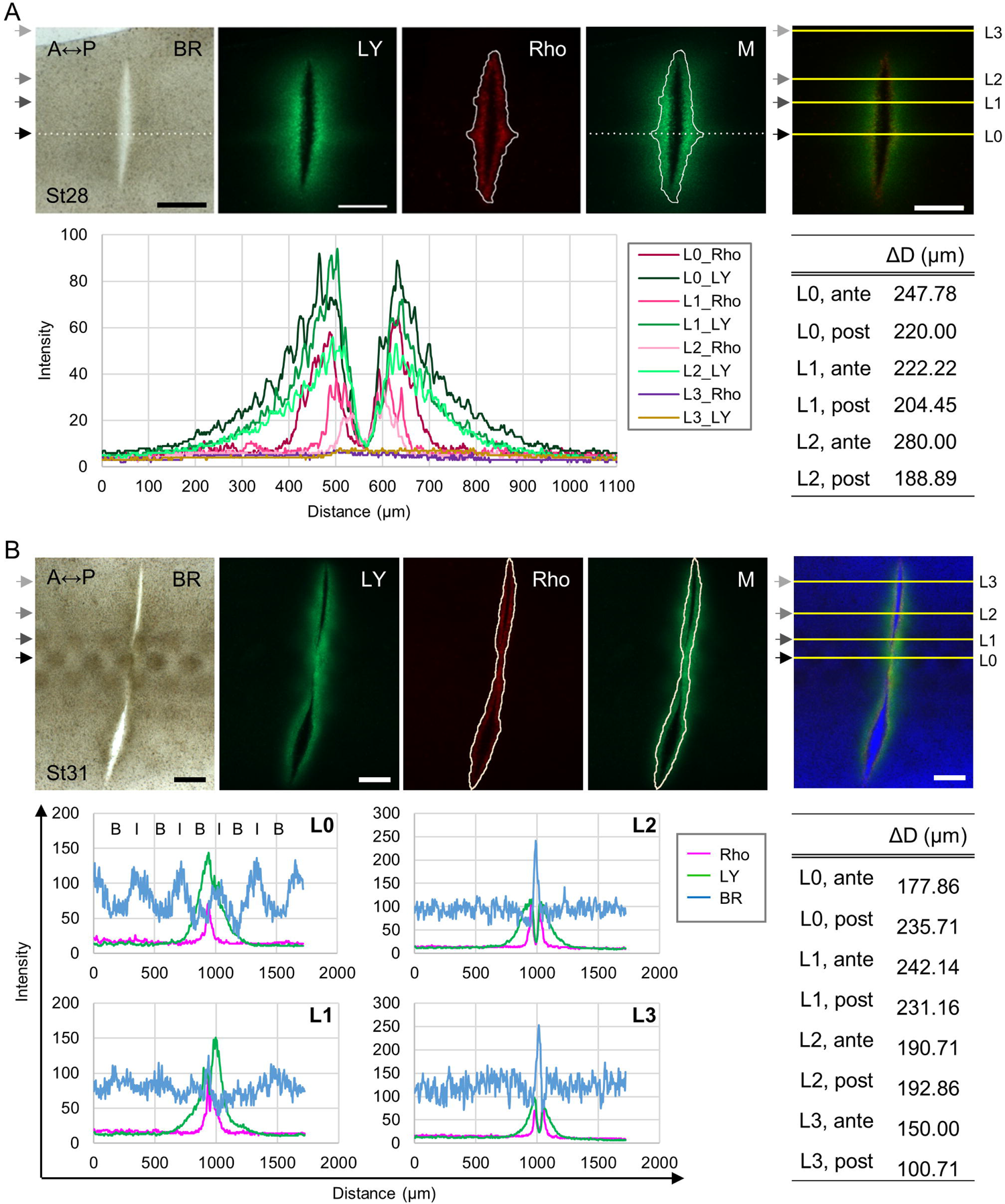
GJIC visualized by scrape-loaded lucifer yellow (LY) dye transfer assay in H&H stage 28-31 chicken dorsal skins. (A) Upper panels: bright-field (BR) and epifluorescent micrographs showing LY dye transfer in the H&H stage 28 skin explant. The dotted lines indicate the midline of the spinal tract. The solid white lines demarcate the edge of the rhodamine dextran (Rho; 10 kDa) signal. Fluorescence of LY (green) and Rhodamine dextran (red) are shown. The merged (M) image was created by overlaying the edge of Rho signal and the midline onto the LY signal. A, anterior. P, posterior. Scale bars, 300 µm. The rightmost panel shows the merged colors of LY and Rho in the ImageJ software. Fluorescence intensities of LY and Rho were measured along the yellow lines and the results were processed by Excel and shown in the lower panels. The triangle-headed arrows in the same color indicate the same position. L, line. Lower panels: distance represents the length of the yellow lines, starting from the anterior. The traveling distance of LY (ΔD) was calculated by the difference in Rho and LY distributions along the yellow lines at the anterior (ante) or the posterior (post) positions. (B) Upper panels: bright-field and epifluorescent micrographs showing LY dye transfer in the H&H stage 31 skin explant. The solid white lines demarcate the edge of the Rho signal. LY is in green. Rhodamine dextran is in red. The merged (M) image was created by overlaying the edge of Rho signal onto the LY signal. A, anterior. P, posterior. Scale bars, 300 µm. The rightmost panel shows the merged colors of BR (converted to blue), LY, and Rho in the ImageJ software. Pixel intensity of BR and fluorescence intensities of LY and Rho were measured along the yellow lines and the results were processed by Excel and shown in the lower panels. The triangle-headed arrows in the same color indicate the same position. L, line. Lower panels: distance represents the length of the yellow lines, starting from the anterior side. The traveling distance of LY (ΔD) was calculated by the difference in Rho and LY distributions along the yellow lines at the anterior (ante) or the posterior (post). B, bud domain. I, interbud domain.

**Figure 7.**
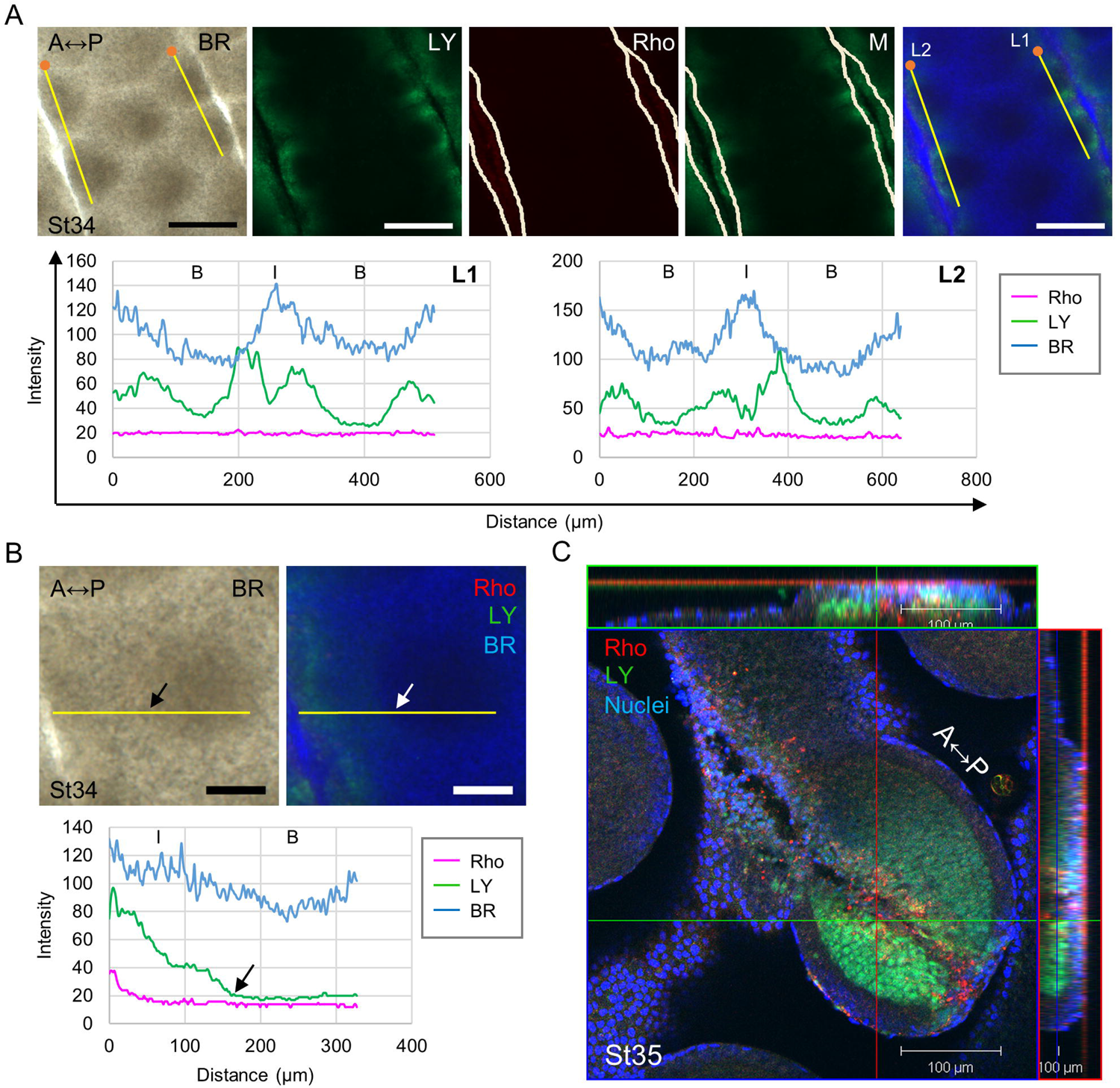
GJIC visualized by scrape-loaded lucifer yellow (LY) dye transfer assay in H&H stage 34-35 chicken dorsal skins. (A) Upper panels: bright-field (BR) and epifluorescent micrographs showing LY dye transfer in the H&H stage 34 skin explant. The solid white lines demarcate the edge of rhodamine dextran (Rho; 10 kDa) signal. LY is in green. Rhodamine dextran is in red. The merged (M) image was created by overlaying the edge of Rho signal onto the LY signal. A, anterior. P, posterior. Scale bars, 300 µm. The rightmost panel shows the merged colors of BR (converted to blue), LY, and Rho in the ImageJ software. Pixel intensity of BR and fluorescence intensities of LY and Rho were measured along the yellow lines starting from the orange dots and the results were processed by Excel and shown in the lower panels. L, line. Lower panels: distance represents the length of the yellow lines. B, bud domain. I, interbud domain. (B) Upper panels: bright-field and epifluorescent micrographs showing LY dye transfer in the H&H stage 34 skin explant. LY is in green. Rho is in red. BR was converted to blue, and the composite image was created in the ImageJ software. A, anterior. P, posterior. Scale bars, 100 µm. Pixel intensity of BR and fluorescence intensities of LY and Rho were measured along the yellow lines and the results were processed by Excel and shown in the lower panel. The triangle-headed arrows indicate the approximate boundary of the bud and interbud domains. Lower panel: distance represents the length of the yellow lines, starting from the anterior. B, bud domain. I, interbud domain. (C) The orthogonal section from z-stack confocal images showing LY dye transfer in a single feather bud and the adjacent area from the H&H stage 35 skin explant. The image was acquired with a Zeiss LSM 510 confocal microscope. LY is in green. Rho is in red. Nuclei were stained by DAPI in blue. A, anterior. P, posterior. Scale bar, 100 µm.

At H&H stage 31, the initial few rows of feather primordia start to form, and the boundary between the bud and interbud becomes visible in the first row (L0) (Figure 6B). The second row of feather primordia (L1) is emerging and is localized at the front of the spreading global patterning wave. L2 and L3 are localized outside the expanding feather tract. The bright-field (BR) image was converted to a blue color using ImageJ software. The lower intensity of the blue signal is associated with the bud domain (B) and the higher intensity is associated with the interbud domain (I). The anterior of the wound along the first row (L0) borders the bud domain and the posterior is adjacent to the interbud domain. We found that dye transfer is slower across the bud domain than the interbud domain in the first row. The space and boundary between the emerging feather primordia in the second row (L1) are less defined, but the cell condensates are visible in the BR image and can be roughly identified by the intensity of the blue signal in the X-Y scatter plot. The anterior wound along the second row (L1) is next to the area with lower cell density and the posterior is adjacent to the higher cell density region. We found that the LY dye traveling distances are similar at both anterior and the posterior regions along the midline (L) and second row (L1) at H&H stage 28. Here the anterior region exhibited a faster dye transfer rate. These results suggest that the interbud region can more efficiently transfer LY dye than the bud in the expanding feather tract. The cells have a similar capacity to communicate with each other when localized at the front of the global patterning wave (H&H stage 31) or when positioned in the emerging feather tract (the midline of the spinal tract; H&H stage 28). The areas localized outside the expanding tract along L2 and L3 exhibited progressively slower LY dye transfer.

At H&H stage 34, short feather buds exhibit apparent A-P polarity and start to elongate. We observed that LY dye transfer is more effective in the interbud region and is gradually reduced towards the center of the bud when the dye was directly loaded into the bud or the interbud region (Figure 7A). We also found that dye transfer was reduced near the center of the interbud region. This indicates that there could be a physical or a molecular barrier between the buds. To support this notion, we showed that the dye loaded into the interbud region could not be transferred into the bud (Figure 7B, arrows indicate the visible boundary of the bud and the interbud region).

At H&H stage 35, the single optical section of the confocal image showed that the elongating feather bud exhibited efficient LY dye coupling in the bud mesoderm (Figure 7C). Dye transfer in the bud epithelium was extremely limited and was not evident in the interbud region. These results suggest that there are physical or molecular barriers between bud epithelium and mesoderm as well as the bud and the interbud region. Taken together, these results showed that GJIC is tightly regulated in both a temporal and spatial manner, and it reflects a dynamic landscape/compartmentalization during early feather morphogenesis, suggesting that GJs facilitate the establishment of boundaries between different compartments.

### Suppression of GJIC activities allows the emergence of new waves of ectopic feather primordia

We next investigated the functional importance of GJIC during early feather morphogenesis by inhibiting GJIC in H&H stage 34 chicken dorsal skin explants with AGA [46–48]. The continuous presence of AGA induced ectopic buds in culture compared to the DMSO control (Figure 8). The bright-field images show the upper thoracic regions (left panels) and the lower thoracic regions (right panels). By day 3 in culture, ectopic buds appeared at the anterior around the base of primary feather buds (arrows) and interbud regions (triangle-headed arrow) (Figure 8A). The size of the ectopic buds became bigger at day 5 (arrows and triangle-headed arrow), and a variety of ectopic bud localizations appeared around the base of primary feather buds (Figure 8B). Interestingly, we rarely observed ectopic buds localized towards the midline around the base of primary feather buds flanking the lower thoracic region. The absence of ectopic buds was highlighted by the dashed circles in magenta. The upper and lower thoracic regions also displayed differential preferences of ectopic bud localizations. The ectopic buds in the upper part tended to appear at both right and left sides around the base of primary feather buds, and the lower part favored the anterior and/or the right or the left side. The observed regional differences suggest that additional factors may influence the capacity of local tissues to respond to the AGA treatment. Some potential candidate factors are cell density and tissue mechanics [9, 49]. Additionally, we observed that the ectopic buds localized in the interbud regions (triangle-headed arrows) appeared at day 2 in culture, about 24 h earlier than the ectopic buds localized at the anterior around the base of primary feather buds (Figure 9A, arrow). To validate the effectiveness of AGA in inhibition of GJIC, we performed a scrape-loaded LY dye transfer assay and observed greatly reduced LY dye transfer upon AGA treatment compared to the DMSO control (A in S2 Figure). GJIC was quantified by measuring the intensity of LY transfer away from the edge of damaged cells (A in S2 Figure, right panel). A non-functional synthetic analog of AGA, glycyrrhizic acid, was used to verify the specificity of AGA, and it does not induce ectopic feather buds (B in S2 Figure). Additionally, we found that carbenoxolone (CBX) [47, 48, 50], the hemisuccinate derivative of AGA, treatment induces ectopic buds similar to the AGA treatment (Figure 9B, arrows and triangle-headed arrows). The ectopic buds express correct patterns of transcriptional markers of feather buds as demonstrated by the whole-mount *in situ* hybridization using the RNA probes targeting *β-catenin* (*β-catenin* expression in the feather tract and developing bud) or *sonic hedgehog* (*Shh*; *Shh* expression at the tip of the bud) (Figure 9C and D) [8, 51–53]. Notably, by day 3 in culture, *β-catenin* expression was detected as a crescent shape at the anterior, extending to the bi-lateral regions around the primary feather buds (Figure 9C). We observed a condensed expression of *β-catenin* at the anterior bud, which is similar to where *Shh* was expressed. This indicates that the anterior has started to form a feather primordium. The expression of *β-catenin* was then restricted to the bud area as the ectopic buds further developed (Figure 9D and C in S2 Figure). Interestingly, we found that evenly spaced ectopic buds, as demonstrated by the *Shh* staining, appeared along the crescent path similar to where *β-catenin* was expressed (Figure 9D, lower panels; arrows and triangle-headed arrows show the ectopic buds at comparable sites before and after staining). These results point to an intriguing possibility that GJIC inhibition stimulates the formation of a new feather tract, and GJ may mediate the exchange of inhibitory factor(s) between neighboring cells to suppress the emergence of ectopic feather buds.

**Figure 8.**
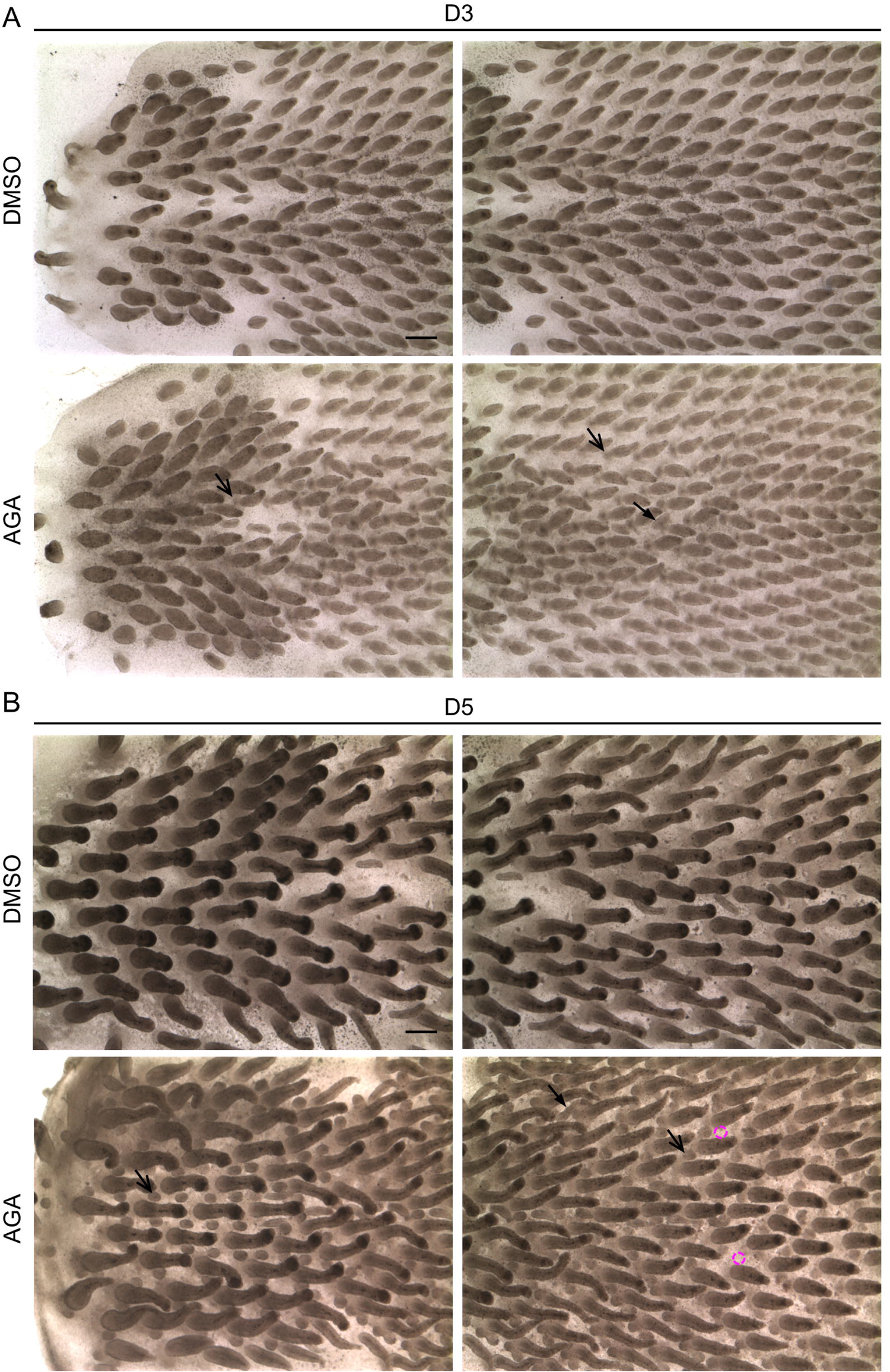
Inhibition of gap junction intercellular communication (GJIC) by 18α-glycyrrhetinic acid (AGA) stimulates the formation of ectopic feather buds. (A-B) Bright-field micrographs showing H&H stage 34 skin explants treated with AGA or DMSO control for three (A) or five (B) days. The arrows indicate the ectopic feather buds localized around the base of the primary feather buds. The triangle-headed arrows show the ectopic buds localized in the interbud regions. Anterior is to the left. Scale bars, 300 µm.

**Figure 9.**
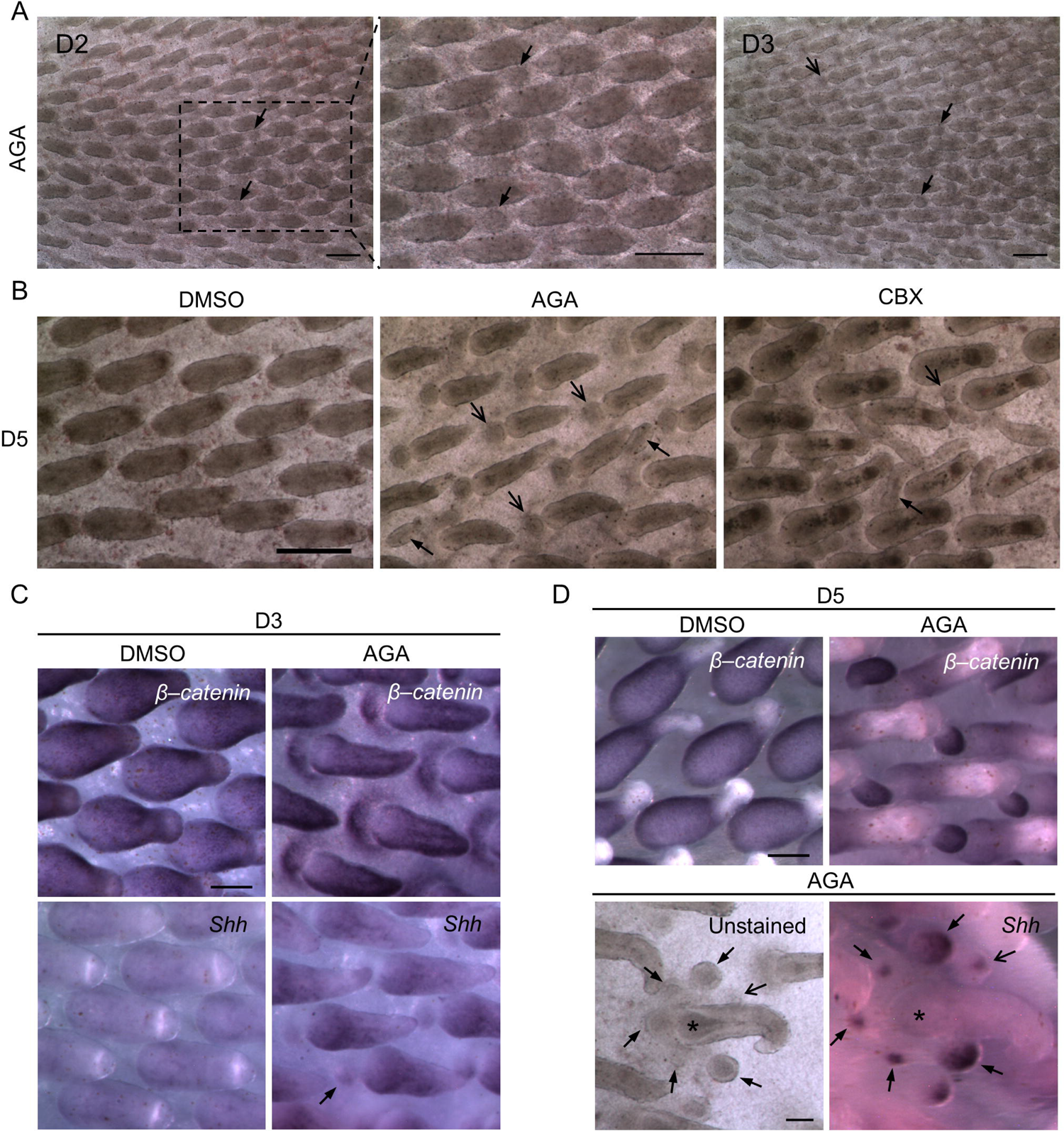
Ectopic feather buds showed molecular markers of feather primordia. (A) Bright-field micrographs showing the H&H stage 34 skin explant treated with AGA for two and three days. The boxed region was enlarged in the middle panel. The triangle-headed arrows indicate the ectopic feather buds localized at comparable sites in the interbud regions. The arrow shows the ectopic bud localized around the base of the primary feather bud. Anterior is to the left. Scale bars, 300 µm. (B) Bright-field micrographs showing H&H stage 34 skin explants treated with DMSO control, AGA or its derivative, carbenoxolone (CBX), for five days. The arrows show the ectopic feather buds localized around the base of the primary feather buds. The triangle-headed arrows indicate the ectopic buds localized in the interbud regions. Anterior is to the left. Scale bar, 300 µm. (C-D) Whole-mount *in situ* hybridization (WM-ISH) of embryonic chicken dorsal skin explants treated with AGA or DMSO control. H&H stage 34 skins were harvested and then *ex vivo* cultured for three (C) or five (D) days. The probes for *in situ* hybridization targeted β-catenin and Shh, the early transcriptional markers of feather primordia formation. Anterior is to the left. Scale bars, 100 µm. (C) The triangle-headed arrow, the expression of Shh at the distal tip of the feather bud. (D) The arrow and triangle-headed arrows indicate ectopic feather buds localized at comparable sites around the primary feather bud (asterisk).

### Exploring the formative process of ectopic feather buds

To further confirm the identity of ectopic feather buds, we performed paraffin-embedded tissue sectioning and H&E staining on the H&H stage 34 skin explants treated with AGA or the DMSO control for three or five days (Figure 10A and B). We focused on the area localized at the anterior side around the base of the primary feather buds because we can unequivocally identify its location on tissue sections. The chicken embryonic epithelium starts to invaginate into the mesoderm at around E11 [54], equivalent to H&H stage 34 (E8) plus three days in culture, and then forms feather follicles. We found that invagination of the epithelium (Figure 10A, triangle-headed arrow) was blocked by the AGA treatment compared to the DMSO control, and this led the epithelium to bulge out (Figure 10A, arrow). This structure keeps growing and starts to show early signs of branching morphogenesis at day five in culture, as shown in the enlarged cross-sectional image (Figure 10B). These histologies show that the epithelial bulged structure exhibits traits of true feather buds and can further differentiate into advanced feather structures. A closer look at the tissue structure around the *in vivo* E10.5-12 chicken embryo dorsal feather tract by scanning electron microscopy confirmed a similar folding/invagination structure observed in the H&H stage 34 skin explants after an additional 3 days of culture (Figure 10C, triangle-headed arrows). These results indicate that the observed developmental processes in the *ex vivo* cultured skin explants have functional relevance to the *in vivo* condition.

**Figure 10.**
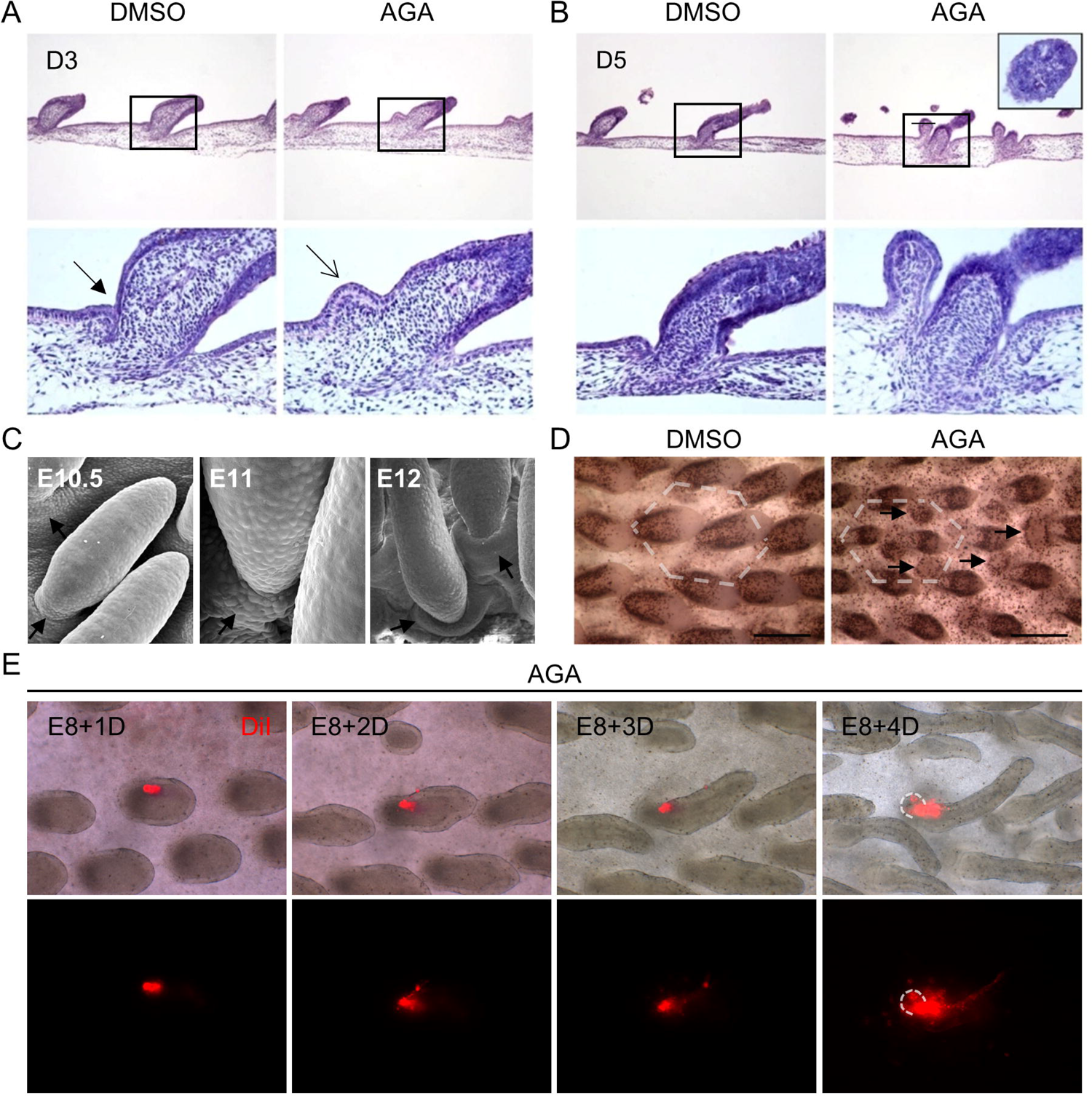
Ectopic feather buds exhibited structural characters of normal feather buds and enhanced cell proliferation. (A-B) Haemotoxylin and eosin (H&E) staining of paraffin-embedded tissue sections obtained from H&H stage 34 skin explants treated with AGA or DMSO control for three or five days. Box regions near the center of the upper images from each day were enlarged and shown in the lower images. The triangle-headed arrow and the arrow indicate similar anatomical locations. The inset within the upper right panel shows an expanded growth along the cross-sectional plane after AGA treatment. Anterior is to the left. (C) Scanning electron microscopic images showing elongated feather buds in the dorsal skins of the E10.5, E11, and E12 chicken embryos. The triangle-headed arrows indicate the folding/invagination structures at the anterior of feather buds. (D) Bright-field micrographs showing the H&H stage 28 dorsal skin explants treated with AGA or the DMSO control for four days and then pulse-labeled with BrdU for four hours followed by immunostaining with the anti-BrdU antibody. Dashed lines indicate the hexagonal feather pattern. The triangle-headed arrows, ectopic feather buds. Anterior is to the left. Scale bars, 300 µm. (E) The H&H stage 34 (E8) skin explant was harvested at day zero and treated with AGA, and then the feather bud was labeled with Vybrant DiI at 25 µM (Invitrogen, in red) at day one of the ex vivo culture. Snap shots of the bright-field and fluorescent images were taken each day after labeling for four days. Dashed lines indicate the ectopic feather bud. Upper panels, merged bright-field and DiI images. Images were processed in the ImageJ software. Lower panels, DiI signals.

Next, we sought to determine whether cell proliferation and movement contribute to the formation of ectopic feather buds. We utilized short-term (4 h) bromodeoxyuridine (BrdU) labeling and whole-mount immunostaining using the antibody against BrdU to visualize proliferating cells in the AGA or DMSO-treated skin explants (Figure 10D). For this experiment, we utilized H&H stage 28 (pre-placode stage) skin explants cultured for four days due to the improved antibody accessibility for whole-mount immunostaining compared to the H&H stage 34 skin explants. We found that the ectopic feather buds exhibit increased cell proliferation upon AGA treatment (Figure 10D, triangle-headed arrows). Additionally, GJIC inhibition does not affect the hexagonal patterning of primary feather buds (Figure 10D, dashed lines). We then used DiI (in red) labeling to track cell movement and found that the cells initially localized in the primary feather bud can contribute to the cell population in the ectopic buds (Figure 10E). These results indicate that the emergence of ectopic feather buds can be attributed to localized cell proliferation and, at least in part, re-localization of cells from the primary feather buds.

A closer molecular characterization of H&H stage 34 skin explants treated with AGA for three days revealed a substantial reduction of Cx43 expression compared to the DMSO control (Figure 11A). We then examined the epithelial tissue structure in H&H stage 34 skin explants treated with AGA or DMSO control for three days by confocal imaging of E-cadherin antibody immunofluorescence (Figure 11B). We found that the basal epithelial cells showed a “rough” membrane structure near the basement membrane upon AGA treatment (Figure 11B, triangle-headed arrows), suggesting an erratic composition of extracellular matrix and communication between basal epithelial cells and dermal cells. Additionally, we observed that there is a transition of basal cell shape from columnar cells (asterisk) to cuboidal cells (hash symbol) when the epithelium starts to invaginate into the mesoderm as shown in the enlarged image from the DMSO control (Figure 11B). In the AGA-treated skin explants, the basal epithelial cells remain columnar in shape (Figure 11B, asterisk in the bottom right panel), suggesting that the epithelium is in a pristine and less differentiated state. Furthermore, we discovered that the expression of transcriptionally active nuclear β-catenin, pY489-β-catenin [[55]], is elevated and accumulates in the ectopic feather bud (Figure 11C). As β-catenin was shown to stimulate ectopic feather formation [8], this result (pY489-β-catenin), together with the observed transcriptional activation of β-catenin from Figure 9C, suggests that AGA treatment re-initiates a developmental program of feather morphogenesis at both post-translational and transcriptional levels.

**Figure 11.**
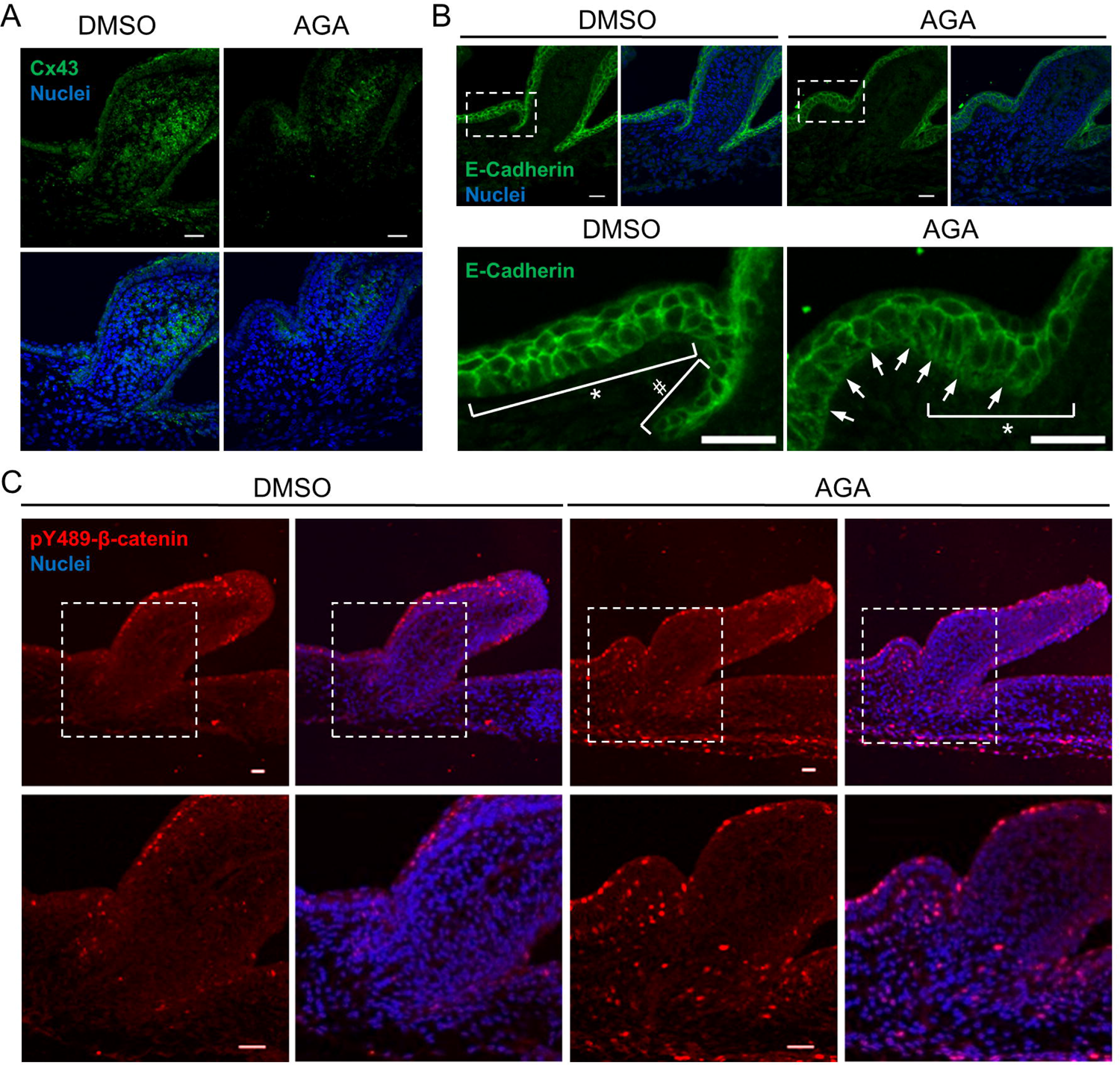
Characterization of ectopic feather buds. (A-C) Immunofluorescence and confocal micrographs showing the expression of the indicated proteins from paraffin-embedded tissue sections of H&H stage 34 skin explants treated with AGA or DMSO (control) for three days. (A) The Cx43 antibody was used for immunostaining (green). Nuclei were stained by DAPI (blue). Scale bars, 20 µm. (B) The antibody against E-Cadherin was used for the immunostaining (green). Nuclei were stained by DAPI (blue). Boxed regions were enlarged below. Asterisks, columnar cells. Pound sign, cuboidal cells. Scale bars 20 µm. (C) The antibody against phosphorylated β–catenin at tyrosine residue 489 (pY489–β–catenin) was used for immunostaining. Boxed regions were enlarged below. Phospho–β–catenin (red). Nuclei were stained by DAPI (blue). Scale bars, 20 µm for the upper panels; 50 µm for the lower panels.

### Mathematical modeling of emerging new buds during feather pattern formation

Our experiments showed that the AGA treatment stimulated multiple waves of ectopic buds. In Figure 12, we summarized the observed patterns at different time points after the AGA treatment in H&H stage 34 skin explants. The ranked order indicates the temporal sequence of primary and ectopic bud appearance. The first wave, patterning the primary buds, is at spot 1. The ectopic buds residing at spot 2 are in the interbud region and show up early between day one and two in culture, thus they grow longer than the ectopic buds in other locations. The additional waves of ectopic buds at other numbered locations start to grow in a crescent-shaped tract at day three in culture and appear over a period of 24 to 48 hours at the anterior and flanked regions around the base of primary buds. The ectopic buds at spot 3 are more frequently observed and show earlier expressions of the feather bud markers (Figure 9C) compared to the spots 4/4’ and 5/5’. The ectopic buds at spot 5/5’ are the least frequently found. The ectopic buds at spot 2 can co-exist with buds at other locations but do not always appear. The observed multiple patterning waves suggested that the AGA treatment can stimulate new instabilities and potentiate the chick skins to initiate a new developmental program until the skins reach a new equilibrium. The first patterning wave acts on a larger scale to set up primary feather buds followed by patterning at smaller scales. Next, we show that such patterns are consistent with a Turing-type model that builds on that developed by Nagorcka and Mooney [56]. Taking inspiration from their work, we use a rectangular domain (specifically, the simulation domain is [0,15] × [0,25]), with periodic boundary conditions top and bottom and zero-flux conditions left and right. Thus, the rectangle represents a small patch of a much larger domain.

**Figure 12.**
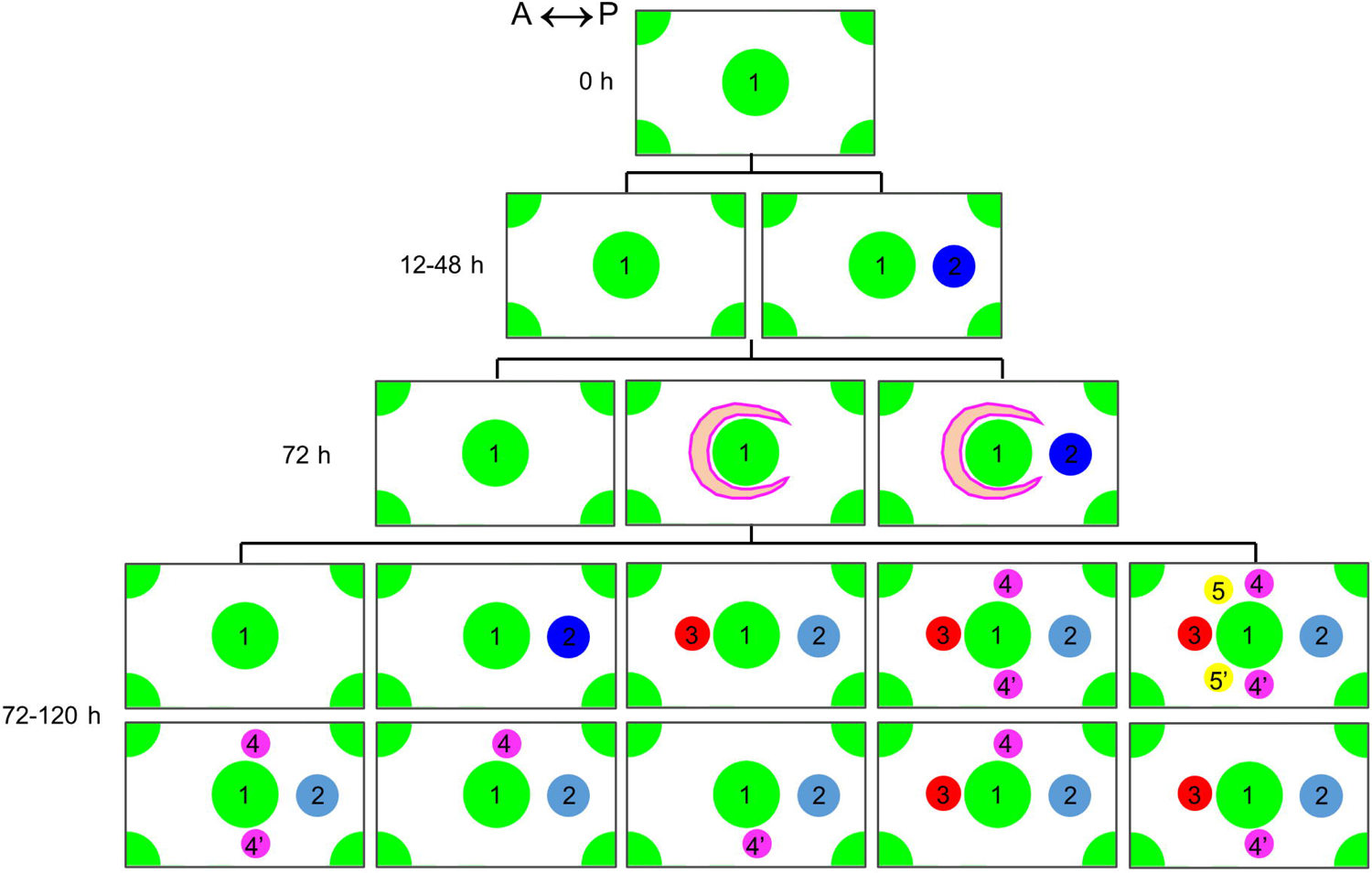
Schematic summary of observed sites of primary and ectopic feather primordia in H&H stage 34 skin explants treated with the GJIC inhibitor. The ranked numbers represent the temporal order of the feather primordia with spot numbered one as the earliest and the primary bud. Spots numbered 2 to 5 represent the ectopic buds. Time 0 h, starting the *ex vivo* culture of H&H stage 34 skin explants. Crescent shapes indicate the emerging field competent to form new buds. Spots in dark or light blue are localized at the same spatial position. The lighter color represents a more flexible state to appear or not to appear in the specific combination of buds as shown in the image. (h, hour). A, anterior. P, posterior.

To produce the spots in the order as numbered in Figure 12, we specify five sets of two coupled diffusing morphogen species, (*u_i_, v_i_*), *i* = 1,2,3,4,5 that can interact in the same way (Figure 13A). Critically, although all species can diffuse throughout the space, interactions are only able to occur in regions that are not already patterned. Namely, at each stage of pattern formation, the tissue through which the morphogens are moving differentiates and does not allow further interactions to occur, again an idea suggested by Nagorcka and Mooney [56]. Specifically, we assume that the tissue undergoes a mechanical change and becomes stiffer, thus, making the morphogens diffuse slower through the tissue.

**Figure 13.**
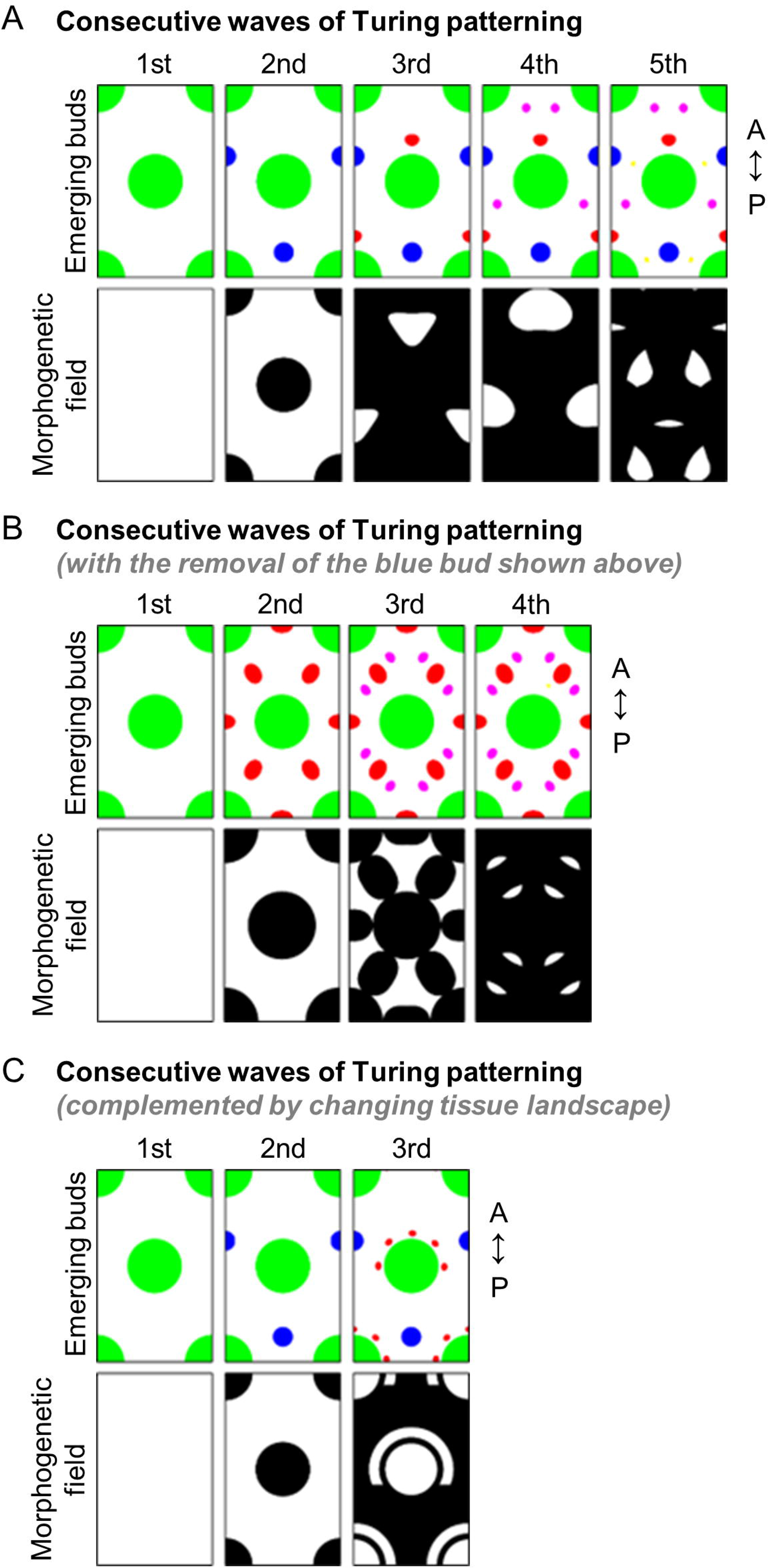
Mathematical modeling of the emergence of ectopic feather buds. (A) Top: Temporal-spatial emergence of ectopic buds. Bottom: Region with competence to undergo patterning is shown in white. Note the dynamic changes of competent regions due to the appearance of buds in the previous stage. All morphogens can diffuse through the domain, however, reactions can only take place in the white region. A, anterior. P, posterior. Animation is provided in S1 Movie. (B) The simulation is the same as that in (A), except spot 2 (blue bud in A) is removed from the simulation sequence and this does not influence the placement of buds 3, 4 and 5. A, anterior. P, posterior. Animation is provided in S2 Movie. (C) Top and bottom represent buds and competent regions as described in (A). Here, we specify the horseshoe region around the first bud in which buds 3, 4 and 5 can appear. A, anterior. P, posterior. Animation is provided in S3 Movie.

Explicitly, the equations underlying the simulations are:

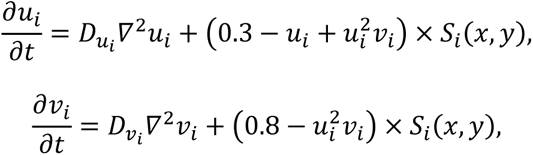

where the kinetics, known generally as Schnakenberg kinetics [57], are fairly arbitrary and are an example of Turing kinetics [1]. Specifically, when the parameters are chosen appropriately, the system will undergo a spontaneous symmetry breaking and produce stationary spatially heterogeneous concentrations of morphogens. These kinetics in particular are well-known for producing simple spot patterns. The *D_ui_* and *D_vi_* are positive constants, which measure the rate of diffusion. Finally, the *S_i_*(*x,y*) function is a piecewise function that only activates the kinetics in specific regions, *Ω_i_*,

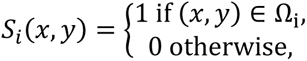

where *Ω_i_* is to be specified in terms of regions not already patterned. A schematic of how these five patterns are formed is shown in the top row of Figure 13A. We start with kinetics that are active everywhere. This system produces the main large spot structure (top left image). Once the first pattern has reached stability, the second Turing system is activated. Critically, although the second set of morphogens is allowed to diffuse everywhere, the reactions are only allowed to occur in the regions where previous activator condensations are not formed. This is shown in Figure 13A. Namely, the second patterning domain is the white region illustrated in the second-from-the-left bottom image. Black regions illustrate where the morphogens can diffuse, but not react. Simulating the system on this domain produces the top second-from-the-left image. This process is repeated until the end of the 5th simulation. In each case of Figure 13A, the bottom image illustrates the morphogenetic field, whilst the corresponding top image illustrates the final pattern that emerges on the given field. Note that spots 3, 4 and 5 require the space to be much more restricted.

Due to the consequential nature of the bud formation, we can perturb the order of bud development. Specifically in Figure 13B we simulate the system again with the same parameter values and the influence of buds 1, 3, 4 and 5 are the same in terms of their spatial restriction. However, we have removed the influence of the second bud (blue bud in Figure 13A). This causes the pattern positioning to appear in unexpected places. This result indicated that the ectopic buds in the interbud region are required to break the anterior-to-posterior positional symmetry of the subsequent patterning waves. Notably, these simulations are solely dependent on coupled activator-inhibitor species from an initially homogenous patterning field. Thus, these simulations may not be able to fully recapitulate the patterning processes in developing embryos, namely the changing tissue landscape. From experimental observations, we notice that buds 3, 4 and 5 seem to be confined to the crescent-shape area and possibly be independent of the placement of bud 2. Therefore, in Figure 13C we simplify the idea of the consecutive simulations by placing a horseshoe region around the first bud to mimic the observed crescent-shaped area around the tissue invagination/evagination site. This specified region is independent of bud 2 and only allows the buds to form in this area. Specifically, the bottom pattern forming field row of Figure 13C illustrates where buds 3, 4 and 5 can appear. In this simulation we showed that local tissue heterogeneity can complement chemical-based Turing’s patterning mechanism to diversify the final patterning outcome. The technical details of our simulations are described in S1 Appendix. The next challenge will be to see if we can relate the morphogens and kinetic interactions to those chemical signals and interactions present in chick feather patterning.

### Connexin 30 is required for the normal periodic formation of primary feather buds in early skin development

We observed that Cx30 exhibits the earliest detectable level of transcripts among the tested connexins as demonstrated by WM-ISH during early feather patterning. Therefore, we further investigated its functional importance by siRNA and replication-competent avian sarcoma virus (RCAS)-mediated overexpression of Cx30 wildtype (a.a. 1-263) and the deletion mutant (a.a. 1-214) lacking the COOH-terminal region (Cx30^ΔC^). We found that perturbation of Cx30 mRNA stability by siRNA through *in ovo* electroporation in embryos at H&H stage 26 (E5) resulted in irregular periodic patterning of the buds (n=6). Random siRNA was used as the control (rc, n=2). Cx30 siRNA treatment resulted in the absence (Figure 14A, asterisk) of feather buds in chick embryos at H&H stage 31 (E7). Some regions show bud fusion (Figure 14A, triangle-headed arrows). We also characterized the impact of siRNA treatment at E5 in older embryos collected at H&H stage 35 (E9) by WM-ISH targeting *Shh* (the feather bud marker expressed in distal bud tip and the marginal plate epithelia of feather filaments) and *Dkk1* (an antagonist of an essential Wnt/β-catenin signaling during feather bud development) (Figure 14B) [58]. A smaller bud with aberrant *Shh* expression appears closer to the anterior indicating mis-orientation of feather bud (arrowhead), and abnormal developing feather filaments (triangle-headed arrow). We also observed dysregulated spacing between buds in the images, although it is not specifically highlighted (Figure 14B). The absence of feather buds in the regions marked by the asterisks did not result from an increase of *Dkk1* expression (Figure 14B). However, this does not exclude the possibility of Dkk1 expression at the protein level or the expression of other antagonists targeting the programs initiating feather development. These results suggested that Cx30 is required for the periodic patterning of feather buds in the early skin development and that Cx30 is involved in the formation of interbud spacing and individual bud development, including size control, orientation, and feather filament morphogenesis.

**Figure 14.**
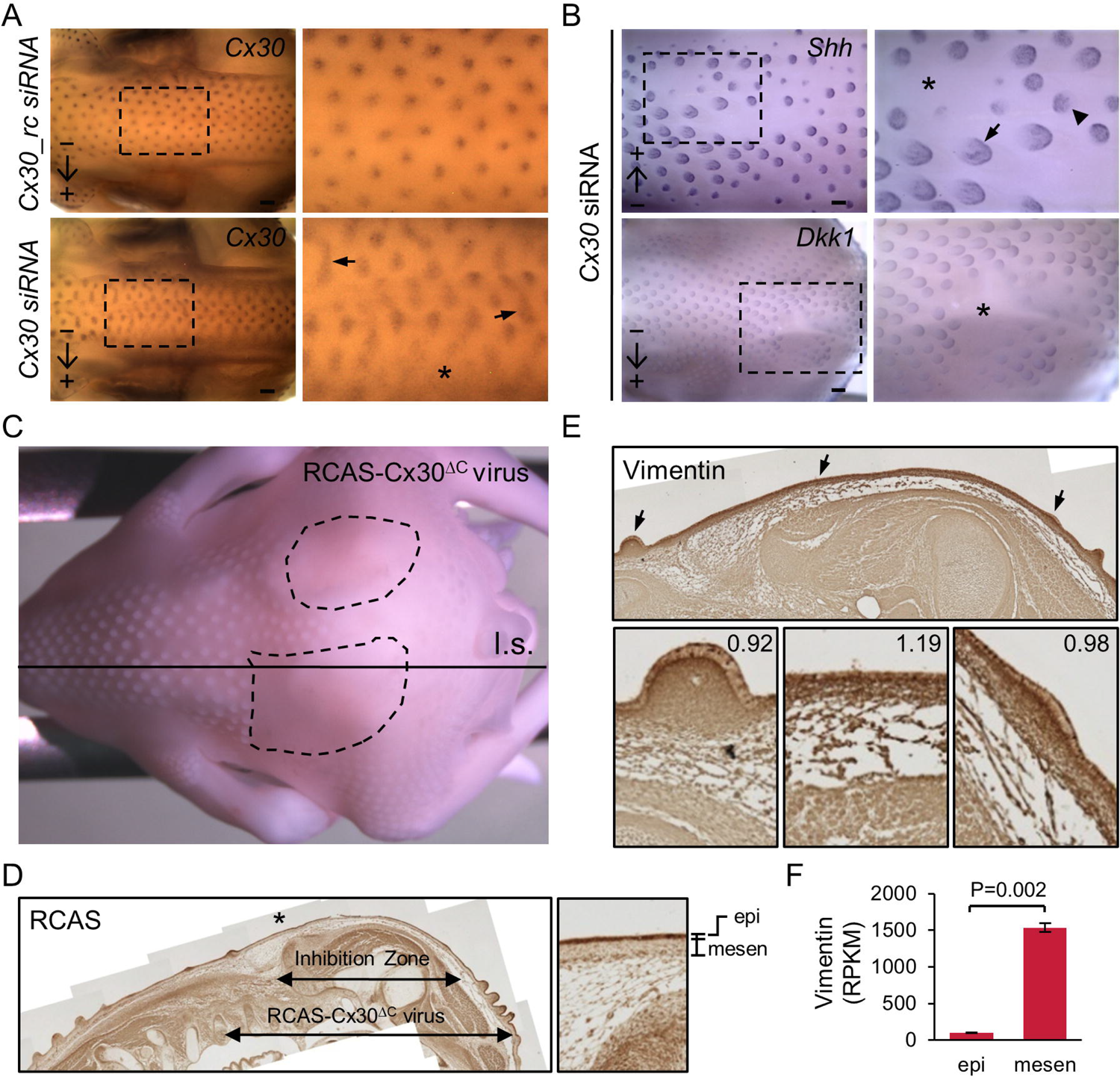
Functional perturbations of connexin 30 by siRNA and a dominant-negative (DN) mutant suppress the formation of periodic feather buds. (A) Cx30 siRNA and the randomized control (rc) were injected and then electroporated *in ovo* into embryos at H&H stage 26. The embryos were collected at H&H stage 31 and stained with Cx30 by WM-ISH. Arrows indicate the direction of siRNA flow. Boxed regions were enlarged on the right panels. The asterisk shows the inhibition zone. The triangle-headed arrows indicate fusions of feather primordia. Scale bars, 300 µm. (B) Cx30 siRNAs were injected and then electroporated *in ovo* into embryos at H&H stage 26. The embryos were collected at H&H stage 35 and stained with probes against *Shh* or *Dkk1* by WM-ISH. Arrows indicate the direction of the siRNA flow. Boxed regions were enlarged on the right panels. The asterisks show the inhibition zones. The triangle-headed arrow indicates abnormal *Shh* expression in differentiating feather filaments. The arrowhead points at the feather bud exhibiting mis-localized *Shh* expression. Scale bars, 300 µm. (C-E) RCAS viruses carrying Cx30 (a.a. 1-214) lacking the COOH-terminal cytosolic region (RCAS-Cx30^ΔC^) were *in ovo* injected into amniotic cavities of embryos at H&H stage 16/17 (E2.5). (C) The RCAS-Cx30^ΔC^ virus-infected embryo was collected at H&H stage 35 and then paraffin-embedded and longitudinally sectioned (l.s.) along the indicated line. The dashed circles mark the inhibition zones. (D) Immunohistochemistry was performed by using the antibody against RCAS viruses and the 3, 3’-diaminobenzidine (DAB) substrate to develop the color (brown). The images of the entire tissue section were manually photo-stitched using Photoshop CS2 software. The double arrows indicate the inhibition zone showing no feather primordia growth and the area with detectable RCAS viruses in the epithelium, respectively. The region near the asterisk was enlarged in the right panel. epi, epithelium. mesen, mesenchyme. (E) Immunohistochemistry was performed in the adjacent tissue section by using the antibody against vimentin and the DAB substrate to develop the color (brown). The areas near the triangle-headed arrows were enlarged in the lower panels. The values indicate the ratio of the vimentin signal in the epithelium to mesenchyme (above the reticular tissue structure) in each image. The intensity of the vimentin signal was calculated by the mean grey value in the manually selected epithelium and mesenchyme areas in the ImageJ software. (F) Reorganization of the previously deposited RNA-seq results (GSE86251) of the gene expression of *vimentin* in the epithelium (epi) or mesenchyme (mesen) of embryos at H&H stage 31. Two biological repeats in each condition. P=0.002 by a two-tailed and unpaired Student’s *t* test.

We next introduced RCAS-Cx30^ΔC^ viruses into the chick embryo at H&H stage 16/17 (E2.5) by *in ovo* injection into the amniotic cavity (Figure 14C, n=2). The RCAS-Cx30^ΔC^ virus-infected embryo exhibited inhibition of feather bud formation at H&H stage 35 (E9) (Figure 14C, the area marked by dashed circles). The presence of RCAS viruses was confirmed by immunohistochemistry (in brown) using the antibody against viral capsid protein p27 in the longitudinally sectioned (l.s.) paraffin-embedded embryo (Figure 14C and D). We found that the RCAS viruses infected mostly the epithelium (highlighted in the image on the right panel, enlarged from the area marked by the asterisk), and the infected area was larger than the zone of inhibition in the tissue section. These results indicate that the RCAS-Cx30^ΔC^ viruses can act dominant-negatively to suppress feather formation, and this process is more effective when a higher concentration of viruses is present in the area. These observations, together with the results of siRNA experiments, confirm the importance of Cx30 in the initiation of feather morphogenesis. We next tried to explore the molecular mechanism of the observed inhibition. Therefore, we performed immunohistochemistry using the antibodies targeting a concise list of candidate proteins in adjacent tissue sections from the RCAS-Cx30^ΔC^ virus-infected embryo (Figure 14D) and found the protein level of Vimentin was elevated in the epithelium of the inhibition zone compared to the nearby regions infected by the viruses (Figure 14E). The images in the lower panels were enlarged from the area indicated by the triangle-headed arrows in the upper panel, and the numbers shown on the upper right corner were obtained from the ratio of the mean intensities of Vimentin in the epithelium versus adjacent dense dermis in the same image. The images were processed using the Image J software. The previous study [38] showed that the RNA or protein level of Vimentin was predominantly expressed in dermal fibroblasts in developing feather buds at H&H stages 31 to 35 (E7-E9). We re-organized their RNA-seq results available from the GEO dataset, GSE86251, for the embryos at H&H stage 31 and present them in Figure 14F. The results that we obtained from the RCAS-Cx30^ΔC^ and Vimentin experiments suggest that Cx30 may facilitate the specification of epithelium during early feather patterning, and this warrants further thorough investigation.

## Discussion

Studies have demonstrated how Turing’s chemical reaction-diffusion (RD) model can be applied to biological systems where morphogen gradient-dependent changes in local cells alter tissue mechanics and generate a global Eda patterning wave that establishes periodic feather bud patterns [2, 9, 10, 15]. Additionally, the zebrafish model showed that cell-cell contact through gap junctions at cell protrusions can modulate skin pigmentation in a Turing-like manner [22]. In the Japanese quail embryonic skin, gap junctions has been shown to be involved in longitudinal color stripe patterning across the body [44], and GJA5 (connexin 40) is involved in stripe and spot formation of intra-feather pigment patterning [59]. Unlike the protein-based morphogens that usually diffuse a short distance, gap junctions can communicate over a long range. Thus, gap junctions serve as good candidates to mediate or modulate the long-range inhibitory signal described in Turing’s RD model. How gap junctions may regulate feather bud patterning is still unknown. In this study, we demonstrate that the expression of GJ isoforms, connexins, and GJIC are highly dynamic and strongly associated with feather morphogenesis, with varied functions at different stages. We further show that GJIC inhibition in the skin explant system by AGA results in uncoupling neighboring cells and allows sequential development of ectopic feather buds at new tissue locations across the embryonic chicken skin. Although the specific molecules going through the gap junction are not fully resolved in this work, our results suggested that GJIC can function as a long-range inhibitory signal during periodic feather patterning. The temporal development of ectopic feather buds can be described by the sequential application of Turing systems that present periodic patterns, as shown in our mathematical model. This work is not in conflict with our original findings that FGF and BMP work as diffusible activators and inhibitors for periodic feather patterning [15] as the potential interplay between different types of morphogenetic signals has not been fully resolved. Yet it reveals an additional layer of GJIC-dependent inhibition in the developing skin. We think the emergence of a new bud is dependent on the net result of interactions between different Turing activator and inhibitor activities, regardless of the forms or sources of those activators / inhibitors.

### Dynamic expression of gap junctions in the developing chicken skin

The permeability of GJs is greatly affected by the composition of GJ isoforms, and different compositions of GJs have their preferred conformation, charge, and substrate size limits [35, 60, 61]. We show 7 different connexin isoforms exhibiting distinct and overlapping temporal and spatial patterns during early feather morphogenesis (summarized in Figure 5), suggesting that GJs can possibly be formed by not only a single GJ isoform but various combinations of GJ isoforms. The complexity of GJ isoform expression in the chicken skin increases as the body grows, and this would allow GJs to modulate more complex cell type-specific behaviors and/or tissue landscapes during development. Importantly, GJIC can also be modulated by protein kinase-mediated post-translational modifications, e.g. phosphorylation [62]. These modifications can rapidly affect the assembly or disassembly of GJs at the plasma membrane. Together with the short half-life of GJ isoforms, around 1-5 hours, these modifications in different types of tissues [63] allow GJs to regulate a dynamic patterning landscape, such as a developing feather field, with messages that can reach long length scales far and fast. Additionally, many of the protein kinases capable of influencing GJIC have known roles in periodic patterning and can be regulated by protein-based morphogens [7, 64, 65]. This raises an interesting possibility that short-range protein-based morphogens and potentially long-range signals mediated by GJs co-function as signaling networks during feather patterning.

We showed that inhibition of GJIC by AGA and its derivative, CBX, can induce ectopic feather buds. This result suggests the existence of inhibitory factor(s) passing through GJs, and this regulation is required to establish correct boundaries and domains during periodic feather patterning. AGA and CBX are more specific and less toxic compared to other GJIC inhibitor types, such as lipophilic molecules (halothane and octanol) and acidifying agents (doxylstearic acids) [47, 50, 66]. AGA was proposed to inhibit GJIC by intercalating into the plasma membrane and binding to GJ hemichannels, leading to closure of GJ channels through conformational changes [47]. CBX was shown to alter arrangements of GJ hemichannels possibly through a direct binding [50]. Notably, we showed in this study that AGA treatment also results in reduced Cx43 protein levels (Figure 11A). Therefore, the ability of Cx43 to act as an adhesion molecule might also contribute to boundary formation and contact inhibition. Because the Cx43 expression and LY dye transfer patterns are similar during early feather patterning, we sought to investigate if specific Cx43 inhibition would lead to a similar phenotype as with AGA treatment. Unfortunately, our attempts to use the specific Cx43 channel blocker, the mimetic peptide GAP27, and siRNA were not successful, likely due to poor tissue permeability and/or short half-life in the skin explant system and embryos (data not shown). Additionally, our previous study showed that lentiviral-mediated Cx43 down-regulation resulted in suppression of feather bud elongation [38]. The lentivirus was injected into embryos at early developmental stages (H&H stage 14 and 17) before feather field formation, and, in the current study, we showed that Cx43 is expressed in the pteric regions at H&H stage 28 (Figure 3). Lentiviral-mediated Cx43 suppression at the early stage might prevent feather field formation, thus blocking future ectopic feathers from forming at a later stage. Whether the effect of AGA is mediated by Cx43, other GJ isoforms, or a combination, will be studied further.

### Gap junction communication may contribute to the Turing long-distance lateral inhibition during feather pattern formation

In this study, we surprisingly uncovered the fact that inhibition of GJIC allows the emergence of four temporal “waves” of new bud formation in *ex vivo* cultured skin explants while maintaining the hexagonal patterning of primary feather buds (Figure 10D). This observation suggests that the primary feather array (spot 1 in Figure 12) is established by the sum of Turing activators / inhibitors, whether they are physical cues or morphogens [9, 67] available at the time of formation. In our experimental conditions that remove the inhibitory activity contributed by gap junctions, new buds emerge in specific locations and “waves”, reflecting the dynamic changes of the morphogenetic competent field (feather tract) once patterning starts. The first wave of ectopic buds emerges in the interbud region with maximal distance to adjacent primary buds (Figure 9A, spot 2 in Figure 12). This suggests that GJIC inhibition may restrict the distance of diffusion of inhibitory factors originated from the primary feather buds. Hence it lowers the threshold for bud growth and allows the emergence of ectopic buds in the initially suppressive interbud region (Figure 12). Second, we found that GJIC inhibition results in increased cell proliferation in the anterior bud base (Figure 10D) and the contribution of cell mass from the primary bud to the ectopic bud (Figure 10E, spot 3 in Figure 12). These results indicate that GJIC inhibition could also allow for the emergence of ectopic feather buds through enhancing the activator signal, hence the cell density [7, 10]. Of note, we found that new buds at spots 4-5 (Figure 12) are localized in a more confined area around the base of the primary feather buds. We speculate that the intrinsic higher cell mass at this region [68] (Figure 10A and C, triangle-headed arrows) provides extra activation cues for the cells in this area to assemble feather buds. Taken together, a hidden layer of Turing-type patterning possibilities is revealed by the removal of gap junctional communications. In our living experimental system, we can explain the results by stating that the threshold for bud formation is reached by the summed activators and inhibitors in that particular space and time.

We then provide a proof-of-concept mathematical model to account for the observed patterning behavior. The classical Turing theory is generally applied to two morphogens acting upon homogeneous environments, which have simple n-dimensional shapes (e.g., lines in one dimension, rectangles in two dimensions, cuboids in three dimensions) with simple boundary conditions. However, even Turing noted that biology presents greater challenges since “most of an organism, most of the time is developing from one pattern into another, rather than from homogeneity into a pattern” [1]. Critically, a developing embryo is a growing 3-dimensional morphogenetic field. As such, tissue curvature and mechanics in developing embryos add extra layers of complexity of engaging diffusive factors in the patterning processes. In our reduced skin explant system and two-dimensional simulations, we minimized these concerns and showed that the Turing-like patterns could emerge under our experimental conditions. Furthermore, our observations of the temporal appearance of stabilized Turing-type patterns in gradually restricted regions following each patterning “wave” suggest the possibility that earlier patterning events could progressively modulate the Turing patterning landscape. Moreover, the self-organizing property of feather buds not only sets up the size and spacing of feather primordia, but also polarity. Preferential accumulation of cells at specific directions or locations, e.g. the clustered progenitor cells around the anterior base of primary feather buds, could reflect tissue mechanics, direction of stress, or flow of morphogenetic cues, and provide activator signal(s) to further modify the patterning landscape [69]. The observed order of temporal patterning is revealed only when we treat skin explants with GJIC inhibitors, thus presumably removing the lateral inhibitors generated by primary feather buds. The appearance of ectopic buds in the interbud regions (spot 2) further showed the possibility of the existence of long-range inhibitor(s) mediated by gap junctions. This is beyond the traditional view of protein-based morphogens which can only diffuse over a much shorter range in a complex tissue system. We suspect the possible long-range inhibitors mediated by gap junctions include the following candidates: ions, e.g., calcium and potassium ions, bioelectricity, second messengers, and tissue mechanics. Notably, in recent years, the identity of factors that are able to generate Turing-like patterns has been greatly expanded in biological systems [2, 10]. It would be exciting to further decipher how biochemical, mechanical, and bioelectrical signals interplay with dynamic cellular behaviors and tissue development, e.g., cell proliferation, migration, differentiation, and body growth, to reach a new stable state in these fast-changing patterning processes.

Research on periodic feather pattern formation has mostly focused on patterning of primary feather follicles. It is still largely unknown how the young and adult feather types are specified and spatially positioned during skin development. Newborn baby chicks are covered by fluffy down feathers which are gradually replaced by adult feather types by weeks 6-12. There are four main forms of feathers that cover the body of the chick, including contour feathers, semiplumes, filoplumes and down feathers. In this work, we did not specify the identity of these emerging buds because skin explants are not suitable for long-term culture. It would be interesting to further investigate how these ectopic buds are correlated with mature feather types in terms of their identity, size, density and spatial localizations. Similar patterning waves have been observed in mammalian hair such as wool secondary follicle formation [70, 71] and fetal hair development [56]. We speculate that GJ-associated patterning mechanisms could be applied to a broader range of living systems during development.

### Suppressive roles of Cx30 at the primary bud formation stage

In developing chicken skin, Cx30 is the earliest detectable transcript, as demonstrated by WM-ISH during feather patterning (Figure 2). We observed that the treatment with Cx30 siRNA or RCAS-Cx30^ΔC^ viruses resulted in inhibition of feather bud formation or fused buds (Figure 14). The observed fusion of feather primordia upon the treatment with Cx30 siRNA (Figure 14A) suggest that the temporal suppression of Cx30 expression may transiently boost the ratio of activator(s) to inhibitor(s), making the ratio lie outside the condition for periodic patterning. The underlying mechanisms will need to be explored further.

We can gain some insights about the cellular and molecular functions of Cx30 from studies conducted in humans and mice. Connexins have been linked to many types of human diseases and congenital skin diseases and they can play diverse functions in different contexts [72–75]. Cx30 (encoded by the *GJB6* gene) mutations in humans account for hearing loss and hidrotic ectodermal dysplasia (Clouston syndrome), which is characterized by severe hair loss, nail hypotrophy, and palmoplantar hyperkeratosis. Human Cx30 (a.a. 1-261) shares 73% of amino acid sequence identity with chick Cx30 (a.a. 1-263). A previous study using human Cx30 (a.a. 1-215), devoid of the COOH-terminal region, showed that this Cx30 mutant mis-localized intracellularly and could not form GJ plaques on the plasma membrane in the HSC-4 human head-and-neck cancer cell line [76]. Cx30 A88V homozygous mutant mouse exhibited leaky Cx30 hemichannels with increased ATP release and calcium ion influx in primary keratinocytes [77], suggesting that GJs formed by wildtype Cx30 could mediate the transfer of ATP and calcium ions. Interestingly, the Cx30 knockout mice created by deletion of the coding region exhibited abnormal endocochlear potential and degeneration of auditory hair cells but, in contrast to humans, showed normal skin and hair development [78]. This is likely due to redundant Cx regulatory networks operating in mouse. Collectively, these studies showed that the embryonic chicken skin has the potential to be used as a model system for studying the contribution of Cx30 in human disease progression. Additionally, it would be of particular interest to know if Cx30 and Cx43 can co-regulate the earliest stage of feather patterning, because these two connexin isoforms both showed early gene expression and the knock-down experiments performed with RCAS-shCx43 in the previous study [38] and siRNA and RCAS-Cx30^ΔC^ viruses in the current work exhibited a similar inhibitory phenotype.

Finally, we qualitatively compared the expression of connexins in developing chicken embryo skin included in this study with what was previously reported [38]. The major aim of this rough comparison is to highlight the connexins expressed in the skin that were not included in this study, and less importantly to provide updated gene names and aliases for each connexin to cross-validate their expression. Of note, these two studies utilized different experimental methods (WM-ISH versus RNA-seq). The RPKM (reads per kilobase million) numbers of the RNA-seq results obtained from the GEO dataset (GSE86251) deposited by the previous study were summarized in S2 Table. The qualitative comparisons were shown in S3 Table. Key observations, excluding the results showing lower expression levels, are: (i) GJB3 and GJC2 are not included in this study but showed moderate and tissue-specific expression in the epithelium and mesenchyme, respectively, in the RNA-seq results; (ii) GJA4 was not detected in this study but showed moderate expression in the mesenchyme and low expression in the epithelium; (iii) GJA1, GJA5, and GJB6 showed moderate to high expressions in both studies; (iv) GJB2 was not included in both studies; (v) most of the connexins showing low expression in one study also exhibited low or no expression in another study. According to these observations, we suggest that future research on early feather patterning should examine the expression and functions of GJB3, GJC2, and GJB2 in more detail.

Overall, the findings shown in this work suggest that gap junctions play a critical role in feather pattern formation. While there are limitations, as mentioned in the discussion above, this study offers new insights into the mechanisms that underlie the emergence of complex spatial patterns in developing skin and in biological systems beyond the skin. A fascinating topic for future investigation is to find out what signaling can be mediated by gap junction communications in periodic patterning, how far and how fast.

## Materials and Methods

### Embryos

Fertilized White Leghorn chicken eggs were obtained from Charles River (Specific pathogen-free) or the local farm (AA Laboratories, Westminster, CA). The eggs were incubated at 38.5 °C in a humidified chamber and staged according to Hamburger and Hamilton (1951).

### Culturing chicken embryonic fibroblasts

Specific pathogen-free St31 chicken embryos were collected and decapitated. Internal organs and limbs were removed with sterile forceps. The embryos were macerated with sterile scissors and then dissociated with 0.05% trypsin with EDTA. The disassociated cells were filtered through 70 µm nylon filter and plated on 100 mm culture dishes in DMEM/high glucose supplemented with 10% fetal calf serum (FCS) and 2% chicken serum.

### Production of RCAS retroviruses

RCAS-Cx30^ΔC^ or the backbone plasmids were transfected into freshly prepared chicken embryonic fibroblasts or DF-1 cells by the treatment of 250 mM CaCl2 and 2X HeBS for 4 hours at 37 °C. Cells were then briefly treated with 15% sterile glycerol in PBS and washed with PBS. Then, they were incubated at 37°C overnight in DMEM with 10% FCS, and 2% CS. The day after transfection, culture media was replaced with 10 mL DMEM with 1% FCS and 0.2% CS. The media was filtered and collected for three consecutive days and then pooled together and centrifuged at 4°C at 20,000 rpm for three hours. One tenth of the original culture media containing viral particles were gently shaken at 4°C overnight. The media were then aliquoted in 100 µL and stored at −80°C until use.

### Injection of RCAS viruses

The aliquoted virus was thawed briefly at 37°C. Volumes of about 5-10 μL were injected into the amniotic cavity of chicken embryos at H&H stage 16/17 (E2.5) or 18 (E3).

### Skin explant culture

The dorsal skins of chicken embryos at St28 or St34 were dissected in Hank’s buffered saline solution under a dissection microscope and then placed onto culture inserts in 6-well culture plates (Falcon, 08-771-15). Skin explants were cultured in DMEM/high glucose supplemented with 10% FBS in a humidified chamber maintained at 37°C at an atmosphere of 5% CO2 and 95% air. The media were replenished every other day.

### GJIC inhibitor treatment

The reversible small molecule inhibitors 18 alpha-glycyrrhetinic acid (Sigma-Aldrich, G8503) and its analog, glycyrrhizic acid (Sigma-Aldrich, 50531), were dissolved in DMSO at a concentration of 50 mM. The reversible AGA-derivative, carbenoxolone (CBX; Sigma-Aldrich, C4790) was dissolved in DPBS at a concentration of 100 mM. The concentrated stock solutions were diluted in DMEM with 10% FBS and 2% CS to make 100 μM working solutions right before the experiments. The culture media were replenished every other day.

### Scrape-loaded lucifer yellow dye transfer assay

Lucifer Yellow CH dipotassium salt (Sigma, L0144) and Rhodamin Dextran (Invitrogen, D1824) were dissolved in distilled water to make a 1% stock solution. The working solution was made by mixing 100 µL lucifer yellow, 100 µL rhodamine dextran stock solution, and 300 µL DPBS. 20-40 µL of working solution was applied to each freshly prepared dorsal skin before scraping, and the LY was allowed to transfer for 8 minutes. Then, the skins were briefly washed with PBS and fixed with 4% paraformaldehyde (PFA) for 20 minutes at room temperature. The samples were imaged by either Zeiss 510 confocal microscope or an epifluorescent microscope.

### Probe making for WM-ISH

The coding sequences of connexins were submitted to Primer3 for primer design. The promoter sequence for T7 RNA polymerase was added on the 5’ end of antisense primer. The primer sequences used in this study were listed in S1 Table. The PCR products at the size of around 500 bp were amplified from the mixture of E7 and E8 chicken cDNAs. The PCR products were then sequenced to verify the identity and transcribed with T7 RNA polymerases to obtain the antisense probes labeled with digoxigenin. The probes for β-catenin and shh were described previously [7].

### siRNA design

The mRNA coding sequence of Cx30 was submitted to Ambion siRNA Target Finder to generate candidate siRNA sequences. The final siRNA sequence was selected according to the following criteria: (i) the sequences having 4 or more Gs in a row were avoided; (ii) the sequences of 30-50% GC were submitted to the BLAST to ensure that they only will anneal to their intended cognate sequence; (iii) no other similarities in the chick genome. The randomized sequence with the same ATCG composition was used as the control. The siRNAs were synthesized by Thermo Scientific. The oligonucleotides for Cx30 siRNA and the randomized control are listed in the S1 Table.

### *In ovo* siRNA electroporation

The siRNAs at the concentration of 100 µM were stored as 10 µL aliquots at −80°C. Immediately before use, a small drop of FastGreen was added to the aliquots, and the siRNAs were used at the highest concentration. The siRNAs were injected into the subepidermal layer of embryos at H&H stage 26 with 1-2 µL. The electrodes were put on the flank of embryos. The siRNAs were transferred to cells using electroporation at 16 V (the actual output is 12 V), 50 ms three times.

### Construction of RCAS plasmids

The COOH-terminal Flag-tagged truncated Cx30 (a.a. 1-214) coding sequence was amplified by the primers listed in S1 Table with the PCR reactions, and then sequenced to make sure the accuracy of the sequence. Then, the target sequence was cloned into the RCAS-2A-mcherry plasmid by ligation. The resulting plasmids were transfected into the One Shot OmniMAX 2-T1R Chemically Competent cells (Invitrogen, C8540-03) by heat shock. The cells containing the desired plasmids were amplified by broth culture and the plasmids were prepared by maxiprep kit (Qiagen, 12663). The plasmids were stored at −20°C until use.

### Sample processing

Embryos for whole-mount *in situ* hybridization (WM-ISH) were collected in DEPC treated phosphate-buffered saline containing 0.1% Tween-20 (DEPC-PBT) and then fixed with 4% PFA in DEPC-treated water at 4°C overnight. Then, samples were dehydrated through a methanol gradient and stored at –20°C. Samples for tissue sectioning were fixed with 4% PFA and then dehydrated through an ethanol gradient. Then, samples were treated with xylene twice and embedded in paraffin. Thickness of tissue sections: 14 μm for WM-ISH samples (except for Cx40, 20 μm) and 7 μm for immunofluorescence and H&E staining.

### Whole-mount *in situ* hybridization (WM-ISH)

Dehydrated embryos were gradually rehydrated into DEPC-PBT and then bleached with 6% H_2_O_2_ in DEPC-PBT followed by treatment with 20 μg/mL Proteinase K in DEPC-PBT. Samples were post-fixed in freshly prepared 0.25% glutaraldehyde/4% PFA solution and then pre-blocked in the hybridization buffer. Digoxigenin-labeled probes were applied to the samples at 65°C overnight. Samples were then washed thoroughly with 2x and 0.2x sodium chloride-sodium phosphate-EDTA Buffer (SSC buffer) for 4 hours. Then, samples were blocked by 20% heat-inactivated goat serum in DEPC-PBT for two hours and treated with pre-absorbed anti-digoxigenin antibody conjugated with alkaline phosphatase at 4°C overnight. Samples were then washed in DEPC-PBT containing 1% levamisole for 3 hours followed by washes in NTMT (100 mM NaCl, 100 mM Tris-HCl, 50 mM MgCl_2_, and 0.1% Tween-20) containing 1% levamisole for 2 hours. Colors were developed with NBT/BCIP, and the reactions were stopped by PBS.

### Immunofluorescence and confocal microscopy

Tissue sections were rehydrated through an ethanol gradient and then treated with 6% H_2_O_2_ in methanol to block the endogenous peroxidase activity. Antigen retrieval was performed with 10 mM citric acid buffer (pH 6.0) at 95°C for 30 minutes. After samples were cooled down to room temperature (RT), they were blocked at RT for two hours followed by incubation with primary antibodies at 4°C overnight. Then, samples were incubated with secondary antibodies conjugated with Alexa Fluor 488 (green) or Alexa Fluor 594 (red) at RT for two hours. After washing, samples were mounted with Vectashield anti–fade medium with DAPI (H-1200, Vector Laboratories). Images were acquired by a Zeiss LSM510 confocal microscope equipped with LSM 510 Version 4.2 SP1 acquisition software (Carl Zeiss). Antibodies: Cx43 (Santa Cruz Biotechnology, sc-9059); E-Cadherin [41]; pY489-β-catenin (Developmental Studies Hybridoma Bank).

### Immunohistochemistry (IHC)

Tissue sections were briefly heated at 65°C for 5 minutes and deparaffinized in xylene, followed by rehydration through an ethanol gradient. Then, 6% H_2_O_2_ in methanol was used to block the endogenous peroxidase. Antigen retrieval was performed with 10 mM citric acid buffer (pH 6.0) at 95°C for 30 min. Sections were allowed to cool down for 30 minutes at room temperature. Then, the sections were pre-blocked by the zeller’s solution for two hours at room temperature. Primary antibodies were diluted in the zeller’s solution and incubated with tissue sections at 4°C overnight. Then, biotin-linked secondary antibodies were applied at 4°C overnight. The next day, streptavidin-horseradish peroxidase was labeled for two hours at room temperature. Colors were developed with AEC (Vector, SK-4200) for 5 to 15 minutes.

### Whole-mount BrdU staining

BrdU stock solution was made by dissolving BrdU powder in Hank’s Buffered Salt Solution (HBSS) at the concentration of 1.5 mg/ml and was stored at −20°C. Skin explants were labeled with 150 µg/ml BrdU for 4 hours and then fixed with 100% methanol for two hours. Then, skin explants were treated with 10% H_2_O_2_ in 1:4 DMSO: Methanol for 2 hours to block the endogenous peroxidase activity. Samples were then treated with 20 μg/mL Proteinase K in PBT at RT for 7 minutes. Then, samples were briefly washed and fixed with 4% paraformaldehyde/ 0.1% glutaraldehyde in PBS at RT for 20 minutes followed by treatment with 2N HCl in PBT for 1 hour. Samples were then treated with 0.1 M sodium borate buffer, pH=8.5 and labeled with the mouse anti-BrdU antibody (Millipore, MAB3424) at 1:1000 dilution followed by the incubation with biotin-linked anti-mouse antibody and streptavidin-horseradish peroxidase. Colors were developed with DAB (Vector, SK-4100).

### DiI labeling

The Vybrant™ DiI cell-labeling solution (Invitrogen, V22885) at the concentration of 25 μM was injected into multiple locations on the skin explants that were harvested at H&H stage 28 and *ex vivo* cultured for one day. Images were taken for four days on an epifluorescent microscope.

### Scanning electron microscopy

Freshly isolated chicken dorsal skins were fixed with Karnovsky’s fixative at 4°C overnight and then post-fixed with thio-carbohydrazide followed by fixation with osmium textroxide. Specimens were critically pointed dried and then coated with gold palladium. Images were acquired with a JEOL JSM-6390LV scanning electron microscope at the Doheny Eye Institute.

### Mathematical simulation

Numerical simulations of the reaction-diffusion systems were run using the finite element software COMSOL Multiphysics 5.3 using a Backward Differentiation Formula scheme in time. The first bud simulation was run for 500 time units before the subsequent reaction-diffusion systems were initiated. Each further reaction-diffusion system was run after 1000 time units. The domain was chosen to have a discretization of 25000 triangular elements. In each case representative simulations were rerun with halved mesh sizes to ensure that the observed patterns did not change with discretization. Additional technical details were listed in S1 Appendix.

**Supporting information** for this article is available online.

## Acknowledgments

We thank Drs. Patricia E. Martin, Eve Kandyba, and Ang Li for helpful discussions. We thank the Cell and Tissue Imaging Core of the USC Research Center for Liver Diseases for imaging studies. This work was supported by National Institutes of Health (NIH) grants R37 AR060306 and RO1 AR078050 and the research contract between USC and China Medical University in Taiwan, contract number 005884. The work was also supported by NIH grant DK048522 to USC Research Center for Liver Diseases. T.E. Woolley and P.K. Maini thank the Mathematical Biosciences Institute (MBI) at Ohio State University for helping to initiate this research. MBI receives its funding through the National Science Foundation grant DMS1440386. We thank the Developmental Studies Hybridoma Bank, created by the NICHD of the NIH, for the maintenance of antibodies at the University of Iowa, Department of Biology, Iowa City, IA 52242.

## Author contributions

CMC conceived the study; CCT and CMC designed the project and experiments; CCT, TXJ, and PW performed the experiments; TEW performed mathematical simulation; PKM, RBW, and CMC provided critical insights and supervised this study; CCT and TEW wrote the manuscript; CCT, TEW, PKM, RBW, and CMC edited the manuscript.

## Supporting information

**S1 Figure. Cx43 expression in the chicken embryo at H&H stage 31.** Cx43 RNA was visualized by whole-mount *in situ* hybridization. Anterior is toward the top.

**S2 Figure. The treatment of AGA effectively suppressed GJIC and induced the formation of ectopic feather buds.** (A) Scrape-loaded LY dye transfer assay. H&H stage 34 skins were harvested and treated with AGA or DMSO control. LY and Rho dyes were loaded the next day. Left two panels: images were taken by the Zeiss LSM 510 confocal microscope. LY is in green. Rho is in red. Anterior is to the left. Scale bars, 100 µm. The right panel shows the fluorescence intensities of LY and Rho along the indicated yellow lines shown in the left two panels. The values of the intensities were obtained using Image J software. (B) Bright-field images showing H&H stage 34 skin explants treated with AGA, glycyrrhizic acid (a non-functional synthetic analog of AGA) or DMSO control for five days. Anterior is to the left. Scale bars, 300 µm. (C) Whole-mount *in situ* hybridization (WM-ISH) of embryonic chicken dorsal skin explants treated with AGA. H&H stage 34 skins were harvested and then *ex vivo* cultured for five days. The probes for *in situ* hybridization targeted β-catenin, the early transcriptional markers of feather primordia formation. Anterior is to the left. Scale bar, 100 µm.

**S1 Table. Oligonucleotides used in the study.**

**S2 Table. The expressions of gap junction isoforms in dorsal skins of chick embryos at H&H stage 31 and 35.**

**S3 Table. Qualitative comparisons of gap junction isoform expressions as shown in this and the earlier study (GSE86251).**

**S1 Data. Raw data used for calculating the traveling distance of lucifer yellow.**

**S1 Movie. Animation of emerging feather primordia.**

**S2 Movie. Animation of emerging feather primordia in the absence of spot 2.**

**S3 Movie. Animation of emerging feather primordia with the complementation of tissue landscape.**

**S1 Appendix. Parameters used in the computational simulations.**

