## Supplementary material for "Gap junctions in Turing-type periodic feather pattern formation": S1 Appendix

**Technical details for the computational simulations**

Definitions of the parameters and active regions during each simulation stage. Note the threshold value defines the morphogen region that becomes feather placode.

Figure 13A

| Stage | $(D_{u_{i}},D_{v_{i}})$ | $\Omega_{i}$ | Threshold |
| --- | --- | --- | --- |
| 1 | (1,40) | $\left[ 0,15 \right]\times[0,25]$, | $u_{Bud}>1.5$ |
| 2 | (3/4,30) | $u_{Bud}<1.5$ | $u_{1}>2.6$ |
| 3 | (2/5,8) | $u_{1}<0.59$ | $u_{2}>2.2$ |
| 4 | (1/10,4) | $u_{2}<0.8$ | $u_{3}>2.4$ |
| 5 | (1/10,4) | $(u_{Bud}>0.8)(u_{1}>0.9)(u_{2}>0.9)$ | $u_{4}>2.55$ |

Figure 13B

| Stage | $(D_{u_{i}},D_{v_{i}})$ | $\Omega_{i}$ | Threshold |
| --- | --- | --- | --- |
| 1 | (1,40) | $\left[ 0,15 \right]\times[0,25]$, | $u_{Bud}>1.5$ |
| 3 | (2/5,8) | $u_{Bud}<1.5$ | $u_{2}>2.2$ |
| 4 | (1/10,4) | $u_{3}<0.8$ | $u_{3}>2.4$ |
| 5 | (1/10,4) | $(u_{Bud}>0.8)(u_{1}>0.9)(u_{2}>0.9)$ | $u_{4}>2.55$ |

Figure 13C

| Stage | $(D_{u_{i}},D_{v_{i}})$ | $\Omega_{i}$ | Threshold |
| --- | --- | --- | --- |
| 1 | (1,40) | $\left[ 0,15 \right]\times[0,25]$, | $u_{Bud}>1.5$ |
| 2 | (3/4,30) | $u_{Bud}<1.5$ | $u_{1}>2.6$ |
| 3, 4, 5 | (1/30,4/3) | $(u_{1}>0.8)(u_{1}<1.2)((y<2)(y>-6)+(y<15)(y>6)+(y<-11))$ | $u_{2}>2.2$ |

**Initial conditions**

For the initial bud simulation the initial condition is a spot of activator in the centre,

$I_{1}\left( x,y \right)=exp(-\left( x-7.5 \right)^{2}-\left( y-12.5 \right)^{2})$. Although a specific condition is used, it is a fairly weak requirement because the same pattern will form under any initial conditions due to the wavelengths chosen by the Turing pattern. The only difference will be that the pattern will be translated left, or right, because of the periodic boundary conditions. Thus, we choose the centered Gaussian to centralize the pattern and, hence, provide simple visual results.

For every other stage the last condition is a source of activator at the back of the bud, $I\left( x,y \right)=\exp\left( -x^{2}-\left( y+10.5 \right)^{2} \right)+\exp\left( -x^{2}-{(y-22.5)}^{2} \right)+exp(-\left( x-15 \right)^{2}-\left( y-22.5 \right)^{2})$.
