## Supplementary figures and images for "Gap junctions in Turing-type periodic feather pattern formation"

### S1 Figure

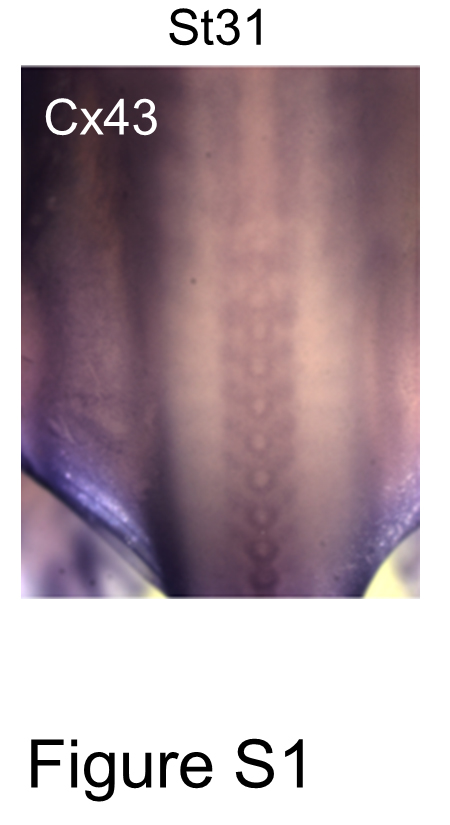

### S2 Figure

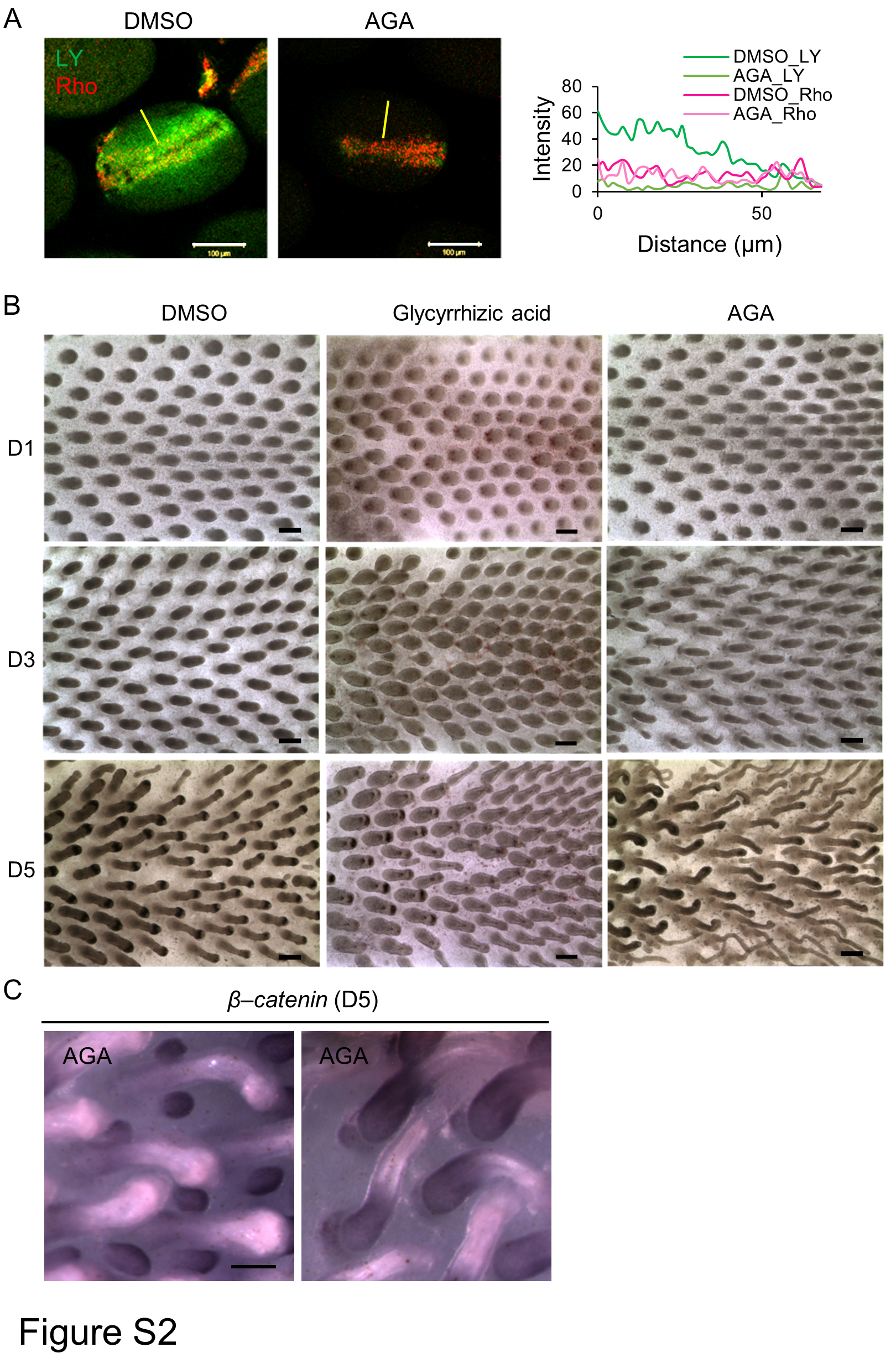
